# Ribosome-Associated Vesicles promote activity-dependent local translation in neurons

**DOI:** 10.1101/2024.06.07.598007

**Authors:** Eva Martin-Solana, Stephen D. Carter, Karthik Krishnamurthy, Giovanni de Nola, Jiying Ning, Jill R. Glausier, Matias A. Preisegger, Laura Hughes, Lauren Meyer, Leanna Eisenman, Paul N. Joseph, Cedric Bouchet-Marquis, Ken Wu, Catherina L. Mobini, Amber N. Frantz, Stephanie Puig, Cheri M. Hampton, Nadine Kabbani, David Mankus, Eric K.F. Donahue, Kristopher Burkewitz, Grant J. Jensen, Simon C. Watkins, Karl Deisseroth, Lief E. Fenno, Michael S. Gold, Zachary P. Wills, Abigail K R Lytton-Jean, Sulagna Das, Zachary Freyberg

## Abstract

Local protein synthesis in axons and dendrites underpins synaptic plasticity. However, the composition of the protein synthesis machinery in distal neuronal processes and the mechanisms for its deployment to local translation sites remain unclear. Here, we employed a multi-scale imaging approach combining cryo-electron tomography, volume electron microscopy, and live-cell imaging to identify endoplasmic reticulum-derived Ribosome-Associated Vesicles (RAVs) as a dynamic platform for moving ribosomes to distal processes and promoting activity-dependent local translation. We demonstrate that neuronal stimulation triggers compartment-specific RAV responses: dendrites accumulate stationary RAVs at sites of enhanced translation, while axons accelerate RAV transport. Real-time imaging of translation at single mRNA resolution reveals that RAVs boost local translation output compared to RAV-independent mechanisms. These findings establish RAVs as specialized platforms that integrate activity-dependent signals with local protein synthesis, providing a mechanistic framework for understanding how neurons achieve precise spatiotemporal control of protein synthesis.

## Introduction

Neurons are highly polarized cells that transfer information through complex networks of synaptic connections. Spatiotemporal control of protein synthesis is essential for neuronal function and plasticity (*1, 2*). Neurons rely on local translation in distal processes (*i.e*., axons and dendrites) to bypass lengthy protein trafficking from the soma (*1, 3-5*). This enables precise control of protein synthesis at distant sites in response to localized stimuli, facilitating time- and site-specific modification of the neuronal proteome (*1*). Disruptions in local protein synthesis underlie neurological and psychiatric disorders, including autism spectrum disorders, fragile X syndrome, and Alzheimer’s disease (*6*). While the importance of local protein synthesis is unequivocal, fundamental questions remain regarding how neurons deliver ribosomes to distal sites and coordinate their activity with local mRNA populations. Electron microscopy studies identified heterogeneous ribosome populations in neuronal processes (*7*), but the molecular mechanisms by which neurons target and coordinate the protein synthesis machinery (*i.e*., mRNAs, ribosomes) to specific sites in the cell periphery remain poorly understood.

mRNAs are directly transported through their association with RNA-binding proteins (RBPs) and trafficked along microtubules (*8, 9*). Recent studies revealed that local translation involves multiple organellar platforms beyond free ribosomes, which are involved in orchestrating mRNA transport, localization, and translation within neurons (*10-16*). For example, endosomes can transport mRNAs and RBPs (*14, 16*), as well as serve as platforms for local translation in neurites (*11*). Mitochondria also associate with ribosomes and co-traffic with nuclear-encoded mitochondrial mRNAs in neurites (*15*). However, despite its well-established role in global protein synthesis, the endoplasmic reticulum (ER) remains relatively understudied in the context of translation outside of the soma, particularly in axons and dendrites.

Traditionally, the ER is divided into rough ER (RER) and smooth ER (SER). RER is characterized by ER cisternae decorated with ribosomes while SER consists of a tubular network that lacks ribosomes. Functionally, the RER serves as a platform for protein synthesis, while the SER is essential for calcium signaling and lipid/cholesterol homeostasis. Notably, though the RER is predominantly found in the soma, extensive RER is absent in axons and is present to only a limited degree in distal dendrites (*15, 17, 18*). Moreover, there is a paucity of ribosomes in the neuronal periphery, with sparse ribosomal populations identified in dendrites and at axonal branch points (*19-21*). Yet, these observations are at odds with a wealth of *in vitro* and *in vivo* biochemical and imaging data that clearly demonstrate local protein synthesis in axons and dendrites (*1, 4, 8, 11, 13, 22-26*). This raises several important questions: how are ribosomes trafficked to distal sites in neurons? Are ribosomes transported to these sites via associations with membranous organelles? Potential answers come from recent evidence of ER-bound ribosomes in axons (*13, 27*), suggesting that the ER may serve as a platform for local translation in the neuron periphery. However, the molecular mechanisms by which the ER and ribosomes engage and regulate protein synthesis outside the cell body remain unclear.

Our previous identification of ribosome-associated vesicles (RAVs) as dynamic RER-derived organelles decorated with 80S ribosomes in various cell types (*28*) prompted us to investigate their potential role in neuronal local translation. We propose that RAVs serve as a platform of local translation to meet the demands for protein synthesis in distal neuronal processes. Here, we employ a multi-scale imaging strategy that integrates live-cell imaging, cryo–electron tomography (cryo-ET), correlative light and electron microscopy (CLEM) under cryogenic and room temperature conditions, and focused ion beam-scanning electron microscopy volume electron microscopy (FIB-SEM VEM) to visualize RAVs in primary neurons and human brain. We demonstrate that RAVs promote local activity-dependent translation in neurites, with distinct behaviors in dendrites and axons. Together, these data provide new insights into the mechanisms underlying local protein synthesis away from the neuronal cell body.

## Results

### Cryo-electron tomography reveals RAVs as ribosome-decorated vesicles in neuronal processes

To establish the presence and ultrastructural characteristics of RAVs in neurons, we conducted correlative light microscopy (CLEM) in rat primary cortical neurons (DIV12) using an antibody that endogenously labels the ER retention signal, KDEL. Consistent with the expected ER distribution in the neurons, KDEL signal was robustly detected in the cell body where the ER is present (Figure 1A). Outside of the cell body, in the neuropil, while we did not see an extensive ER network, we observed KDEL-positive signal that colocalized with vesicular structures (246 nm ± 99.8, n=14) (Figure 1B), consistent with the dimensions of RAVs (*28*). To further characterize these vesicular structures *in situ* under near-native conditions, we conducted cryo-correlative light and electron microscopy (cryo-CLEM) in primary neurons expressing KDEL tagged with mNeonGreen (mNeon-KDEL) (DIV 10-12) (Figure 1C-G, S1). Consistent with RAVs, Cryo-CLEM confirmed that mNeon-KDEL puncta localized to vesicular structures decorated with electron-dense ribosome-like particles (Figure 1C-D, Movie S1). Among the 36 acquired tomograms, 42% of the structures that co-localized with mNeon-KDEL were identified as RAVs (Figure S1A, S1C), while 28% corresponded to ribosome-free vesicles (Figure S1B, S1C). The average RAV diameter was ∼400 nm (395.9 ± 140.5 nm, n=19) consistent with previous reports (*28*), while ribosome-free vesicles were slightly smaller (215.5 ± 107 nm, n=19) (Figure S1D). To corroborate that the electron-dense particles associated with RAV membranes were ribosomes, we performed subtomogram averaging (STA) of the particles (n=164). STA resolved 80S mammalian ribosomes at ∼40 Å resolution (Figure 1E, S1E and S1F). Moreover, we confirmed that the three-dimensional (3D) structure of the averaged particles strongly resembled a well-defined 80S ribosome structure from *Oryctolagus cuniculus* previously imaged by cryo-ET (Electron Microscopy Data Bank #EMD-0529) that was down-sampled to 32 Å resolution (*29*) (Figure 1F). Mapping the ribosomes back to their original positions revealed that the ribosomes decorated the RAV membranes (Figure 1G).

**Figure 1.**
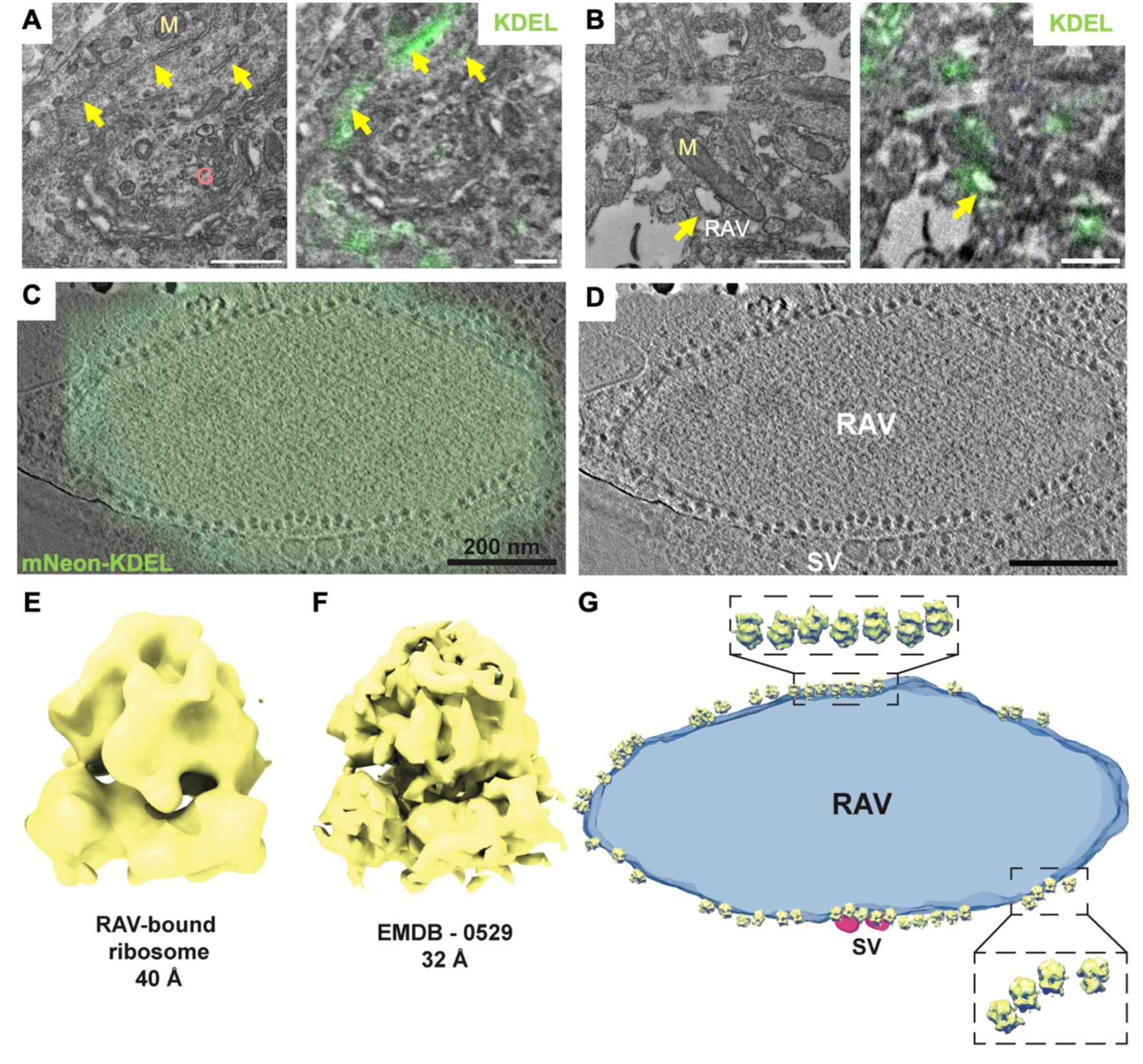
Identification of RAVs by correlative light and electron microscopy (CLEM) of KDEL-labeled structures. **(A, B)** CLEM imaging of endoplasmic reticulum (ER) distribution in primary rat cortical neurons using an anti-KDEL antibody. **(A)** A representative high-magnification micrograph (left panel) demonstrated extensive ER network in the cell soma. Correlative image (right panel) showing colocalization of anti-KDEL signal (in green) with ER (arrowhead). The Golgi apparatus (G) and mitochondria (M) are also visible. Scale bar, 1 µm. **(B)** Higher magnification of the neuropil regions showing RAVs. Right panel: correlative images showing the same vesicular structures colocalizing with anti-KDEL antibody. Scale bar, 500 nm. **(C, D)** Cryo-CLEM shows ER marker mNeon-KDEL localized to discrete puncta in the periphery of primary hippocampal neurons. A cryo-tomographic slice with **(C)** and without **(D)** overlay of an epifluorescence image with mNeon-KDEL fluorescence (in green). Scale bars, 200 nm. **(E)** Average of 164 manually selected subvolumes of RAV-bound particles from 5 tomograms. Reference-free alignment provided a structure consistent with an 80S mammalian ribosome. **(F)** Subtomogram average of a validated mammalian 80S ribosome structure (EMD-0529; down-sampled to 32Å) by cryo-ET, indicating RAV-bound subvolumes were 80S ribosomes. **(G)** Isosurface of the RAV in Panel C featuring subtomogram averaged RAV-bound 80S ribosomes mapped back to their original positions. Enlarged views indicated an ordered polysome arrangement of RAV-associated ribosomes.

### Identification of RAVs in human brain tissue

We next examined whether RAVs are also present within intact tissue, specifically human brain. We therefore imaged human cortex via FIB-SEM VEM. First, we confirmed that the overall ultrastructural organization was well preserved as evidenced by the presence of intact organelles such as mitochondria and the Golgi apparatus within the neuronal cell body and neuropil (Figure S2A, Movie S2). We also observed an extensive network of RER in the soma, characterized by membrane sheets with associated ribosome-like electron-dense particles (Figure S2A, S2B). Notably, we identified vesicular structures in the cell body with membrane-associated ribosome-like densities that resembled RER, consistent with putative RAVs (Figure S2C, S2D). Indeed, measurements of the membrane-bound ribosome-like particles from the segmented RAV yielded a mean diameter of 33.6 ± 4.6 nm (n = 25), consistent with the diameter of mammalian 80S ribosomes (∼30 nm) (Figure S2D, S2E, Movie S3). Similarly, we observed RAV-like structures in the neuropil that resembled those in the cell body (Figure S2F, S2G). Although RAV-like vesicles in the cell body had a larger diameter (418.7 ± 219.0 nm, n=30) compared to those in the neuropil (264.9 ± 65.4 nm, n=9) (Figure S2H), the overall diameters of all RAV-like vesicles in human brain tissue were consistent with those previously reported (*28*).

### RAVs exhibit dynamic microtubule-dependent trafficking in neuronal processes

We next examined the dynamics of RAV trafficking in primary neurons by live-cell wide-field microscopy. We previously used different ER markers to label RAVs in pancreatic β-cells, strongly suggesting that RAVs are a dynamic subcompartment of the ER (*28*). Here, in line with our earlier work, we observed dynamic, well-defined punctate structures that were labeled using various ER markers [*i.e*., mNeon-KDEL (*28*), ER-mScarlet (*30*), Sec61β-GFP (*28*)] in primary neurons (Figure S3). ER-mScarlet exhibited the strongest, more consistent signal, justifying its choice as a marker for RAV visualization. Using this ER marker, we identified two subpopulations of RAVs in primary cortical neurons: actively moving versus stationary RAVs. The majority of RAVs were moving (66% of total RAVs), while the remainder were stationary (34%) (Figure 2A-D, Movie S4). Kymograph analysis of the moving RAVs revealed that 59% of RAVs underwent processive movement, whereas the rest (41%) displayed non-continuous or intermittent motion characterized by pausing for variable time periods along their paths (Figure 2D). Moreover, moving RAVs showed a wide distribution of speeds ranging from 0.2-2.8 µm/s (Figure 2B. 2F) that was bidirectional (Figure 2A-C, Movie S4), consistent with microtubule-mediated transport (*31, 32*). To confirm that RAV trafficking in neurons relies on microtubule-mediated transport, we analyzed the impact of microtubule disruption via nocodazole treatment (1µM, 2h) on RAV movement in rat hippocampal neurons. Nocodazole is a known antimitotic agent that destabilizes the microtubule network by inhibiting microtubule polymerization (*33*). Following treatment, the proportion of processive RAVs significantly decreased [control, processive: 202/446 (45%); nocodazole, processive: 78/362 (22%); p <0.0001]. In contrast, the proportion of stationary RAVs significantly increased [control, stationary: 117/446 (26%); (45%); nocodazole, stationary:182/362 (22%); p = 0.0290] (Figure 2E). Interestingly, there was no change in the proportion of intermittently processive RAVs [control, intermittent=129/446 (29%); nocodazole, intermittent: 102/262 (28%); p = 0.1924] (Figure 2E). Furthermore, the speed of actively moving RAVs decreased 1.8-fold following nocodazole treatment (control 1.08 ± 0.05 µm/s; nocodazole 0.59 ± 0.05) (Figure 2F). Taken together, these results suggest that RAV trafficking is mediated by microtubules, consistent with previous studies showing that the ER uses motor proteins (*i.e*., kinesin and dynein) for its transport (*34-36*).

**Figure 2.**
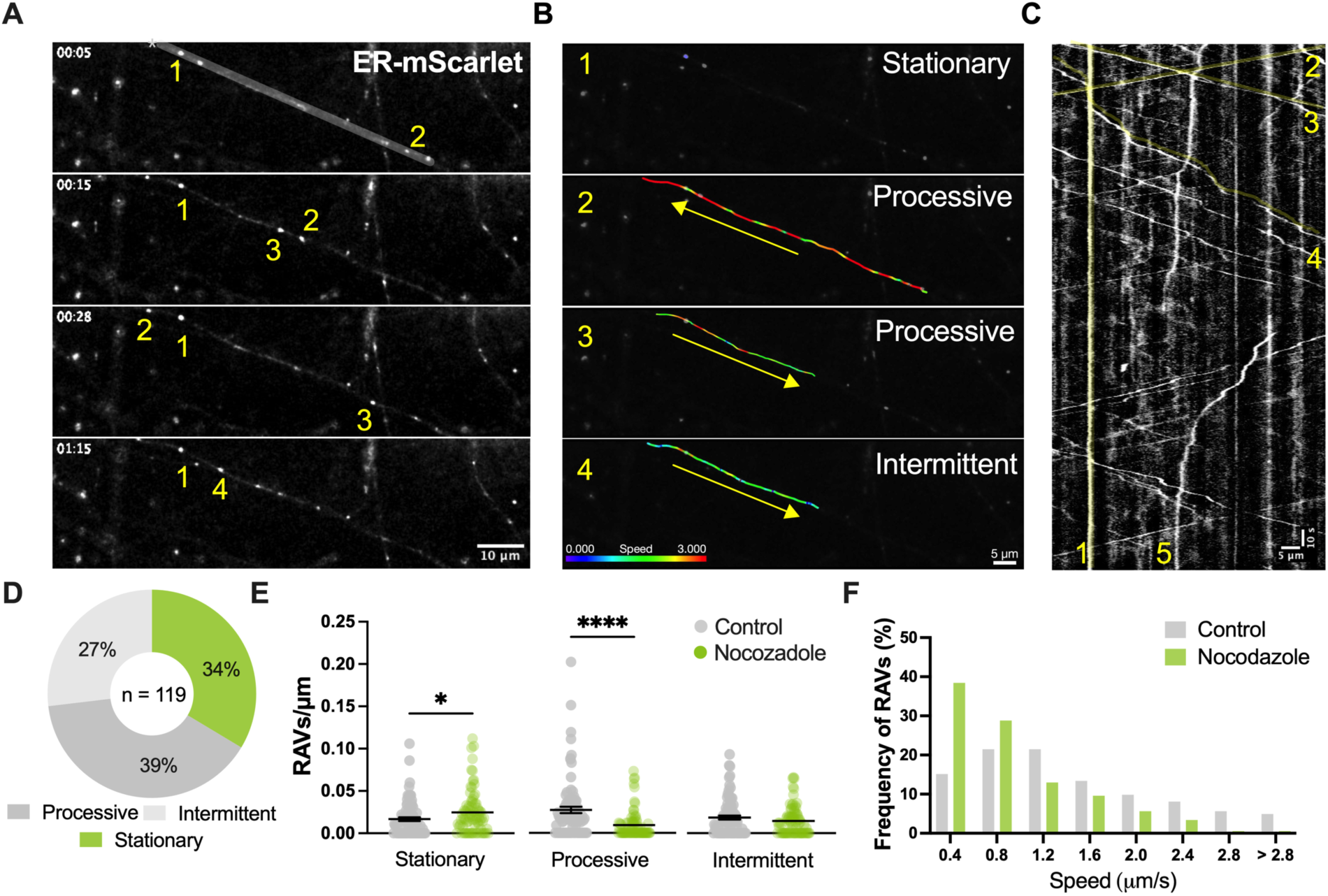
RAV dynamics in neurites/distal neurites periphery. **(A)** Representative time-lapse images demonstrate the dynamics of ER-mScarletI-labeled RAVs (numbered 1-4) over time along a neurite of a primary cortical neuron (DIV11-12). Scale bar, 10 μm. **(B)** Frame-by-frame velocity measurements of RAVs in Panel A. Each of the 4 highlighted RAVs (1-4) demonstrated different dynamics as indicated by the color-coded velocity range: stationary (1), fast (2), moderate (3), and slow (4) velocities, respectively. Arrows specify the overall direction of movement along the neurite. Scale bar, 5 μm. **(C)** Kymograph of the segmented neurite from Panel A. Highlighted lines (in yellow) indicate movements of the RAVs (1-4) featured in Panels A and B (1-4) across a 5-min timespan. (5) indicates a RAV with intermittent movement prior to stopping in place. Scale bar, 5 μm, 10 sec imaging. **(D)** Relative proportion of moving (processive and intermittent) versus stationary RAVs in the periphery (n=119, 30 neurites). **(E)** RAV density after nocodazole treatment (1 µm, 2 h) (DIV12). Mean ± SEM; *p<0.05, **p < 0.01, ***p < 0.001, ****p < 0.0001, Unpaired t-test. **(F)** Distribution of average speeds of moving RAVs in response to nocodazole treatment.

### Neuronal activity triggers compartment-specific RAV dynamics

Substantial evidence demonstrates enhanced local protein synthesis in distal neuronal processes which contributes to activity-dependent plasticity (*25, 37-39*). We therefore determined whether neuronal stimulation alters RAV trafficking using an established chemically-induced long-term potentiation (cLTP) paradigm (*40*). In response to cLTP stimulation of primary cortical neurons, the density of RAVs in neurites increased significantly by 1.5-fold within 15 min of cLTP (0.032 ± 0.003 RAVs/µm) compared to basal conditions prior to cLTP (0.022 ± 0.003 RAVs/µm) (p = 0.0418) (Figure S4A, Table 1). The increase in RAV density persisted for 60 min after cLTP (0.031 ± 0.004 RAVs/µm) (p = 0.2427) (Figure S4A, Table 1). However, cLTP did not modify the average speed of moving RAVs in the periphery (Table 1). Given the absence of significant differences between 15 and 60 min of cLTP, we focused primarily on the earlier time point for subsequent analyses.

We next validated our results in primary hippocampal neurons, which also exhibited an increase in RAV numbers after 15 min of cLTP compared to pre-stimulation [cLTP: 0.085 ± 0.005; pre-cLTP: 0.071 ± 0.005; p = 0.0405] (Figure S4B, Table 2). Indeed, we detected significantly more RAV-containing neurites in response to cLTP (12.9 ± 4.3% increase, p = 0.028) (Figure S4B). This activity-driven increase was dominated by increased localization of RAVs to dendrites, where cLTP stimulation was associated with ∼60% increase in RAV density (cLTP: 0.099 ± 0.011; pre-cLTP: 0.062 ± 0.005 RAVs/µm; p = 0.050) (Figure 3A-E). The increase in dendritic RAVs was primarily due to an increase in stationary RAVs (cLTP: 0.067± 0.012; pre-cLTP: 0.040 ± 0.005 RAVs/µm; p = 0.0113). There were no accompanying changes in moving RAVs within distal dendrites (cLTP: 0.032 ± 0.084; pre-cLTP: 0.022 ± 0.006 RAVs/µm; p = 0.4421) (Figure 3F, 3G, Table S3). These data suggest that previously moving RAVs from elsewhere in the neuron enter dendrites and become stationary within this initial window of cell stimulation.

**Figure 3.**
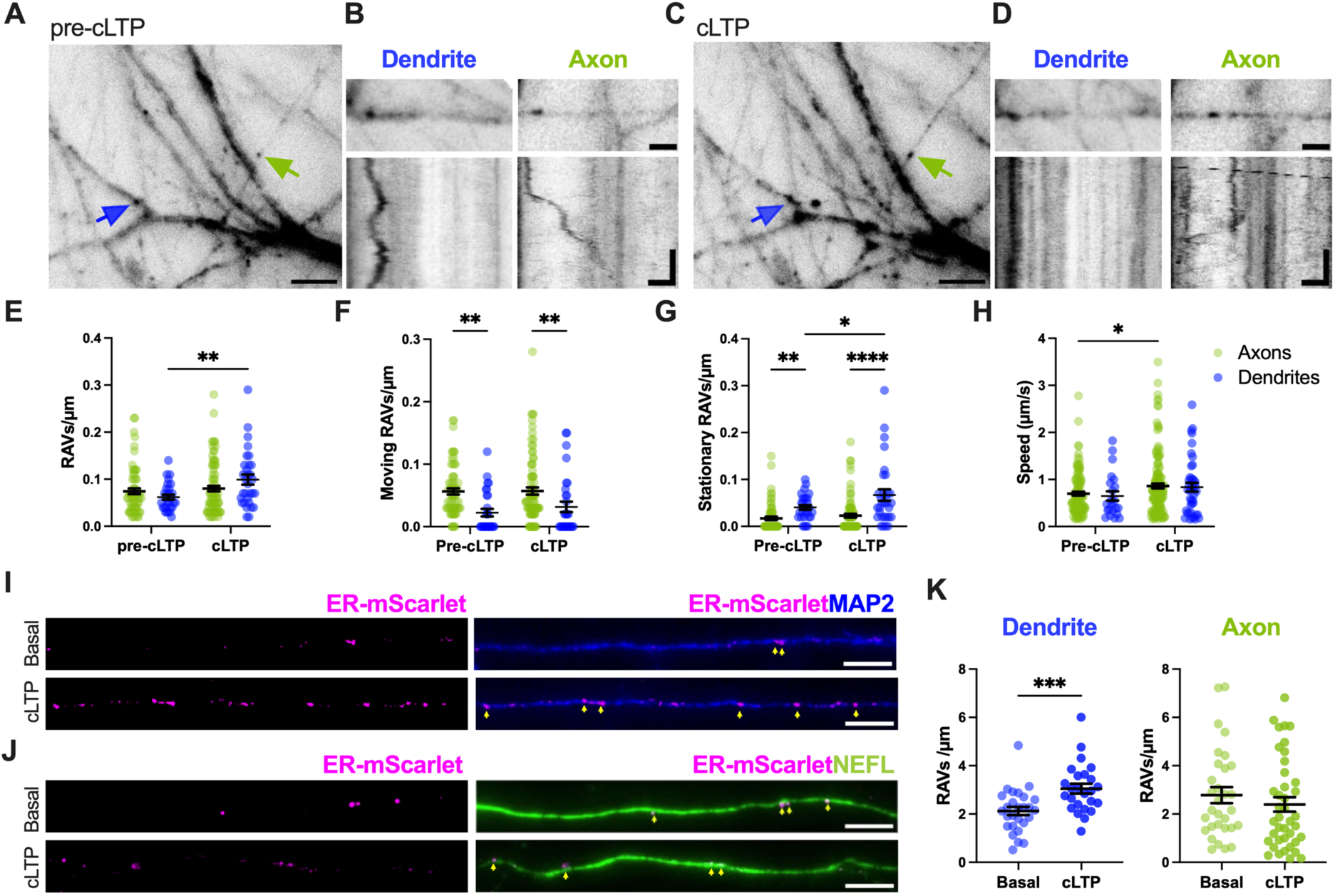
cLTP-driven changes in RAV dynamics differ in axons versus dendrites. **(A-D)** Analysis of RAV dynamics in mouse primary hippocampal neurons. **(A and C)**. Representative image pre- **(A)**, and post-cLTP **(C)** showing RAV trafficking in dendrites and axons. Scale bar, 5 µm. (B and D) Straightened dendrite and axons pre- **(B)** and post-cLTP **(D)** as indicated by the respective arrows from Panels A and C, and their representative kymographs comparing RAV movement in axons versus dendrites. Scale bar, 2 µm. Scale bar kymographs, 2 µm, 20 sec **(E-H).** Quantitative analyses of RAV dynamics in hippocampal axons and dendrites in response to cLTP including: total RAV density **(E)**, density of moving RAVs **(F)**, density of stationary RAVs **(G)**, and RAV speed **(H)**; Two-way ANOVA, Fisher’s LSD test. n= 66 axons (pre-cLTP), n= 29 dendrites (pre-cLTP), n = 87 axons (cLTP) and n = 31 dendrites (cLTP). **(I, J)** Representative immunofluorescence images confirming RAV localization to MAP2+ dendrites **(I)** and NEFL+ axons **(J)** under basal pre-cLTP and cLTP conditions. Scale bar, 5 µM. **(K)** Quantification of immunofluorescence data showed RAV density in dendrites and axons under basal versus cLTP conditions. Unpaired t-test. n = 40 axons (pre-cLTP), n = 27 dendrites (pre-cLTP), n = 31 axons (cLTP), n= 25 dendrites (cLTP). Mean ± SEM; *p<0.05, **p < 0.01, ***p < 0.001, ****p < 0.0001.

Live-cell imaging also revealed numerous RAVs moving in thin, spineless processes distal from the cell body, consistent with axons (Figure 3A-D). However, unlike dendrites, there was no cLTP-induced change either in the total density of axonal RAVs (cLTP: 0.080 ± 0.006 RAVs/µm; pre-cLTP: 0.074 ± 0.061 RAVs/µm; p = 0.4922), or in the density of moving (cLTP: 0.057 ± 0.006; pre-cLTP: 0.056 ± 0.050 RAVs/µm; p = 0.9317) and stationary axonal RAVs (cLTP: 0.023 ± 0.004; pre-cLTP: 0.017 ± 0.004 RAVs/µm; p = 0.3797) (Figure 3E-G, Table S3). Furthermore, cLTP caused an increase in RAV speed in axons (cLTP: 0.086 ± 0.05 RAVs/µm; pre-cLTP: 0.70 ± 0.04 RAVs/µm; p = 0.0175), but not in dendrites (cLTP: 0.84 ± 0.09 RAVs/µm; pre-cLTP: 0.65 ± 0.10 RAVs/µm; p = 0.1995) (Figure 3H, Table S3). Notably, we found that RAVs were more stationary in dendrites (cLTP: 0.067± 0.012; pre-cLTP: 0.040 ± 0.005 RAVs/µm) compared to axons (cLTP: 0.023 ± 0.004; pre-cLTP: 0.017 ± 0.004 RAVs/µm) (pre-cLTP, p = 0.0099; cLTP; p < 0.0001) (Figure 3G). Additionally, we identified RAVs in mouse dorsal root ganglion neurons (DRGs) which exclusively possess axons but not dendrites (Figure S4C, Movie S5).

Finally, to further confirm the presence of RAV in axons and dendrites, we immunolabelled these processes with established markers [*i.e*., MAP2 (Microtubule-Associated Protein 2) for dendrites, NEFL (Neurofilament Light Chain) for axons] (Figure 3I, 3J). Quantification of RAVs in the two distinct processes was consistent with our previous data, showing that RAV density significantly increased in dendrites after cLTP (cLTP: 3.05 ± 0.20; basal: 2.12 ± 0.17; p = 0.0010) while remaining unchanged in axons (cLTP: 2.39 ± 0.30; basal: 2.78 ± 0.33; p = 0.3871) (Figure 3K). Overall, these data suggest that RAV trafficking in axons and dendrites is sensitive to neuronal stimulation, albeit in ways distinct to each subcompartment.

### RAVs serve as platforms for activity-dependent local translation

Local translation in the neuron periphery requires spatiotemporal coordination of both mRNA and ribosome trafficking (*1, 5*). However, many questions remain concerning the precise relationships between mRNAs and the ribosomal machinery in axons and dendrites. We therefore examined co-trafficking between endogenous β-actin mRNA, which is known to be trafficked and translated in an activity-dependent manner (*25, 41, 42*), alongside oScarlet-KDEL-labeled-RAVs in primary hippocampal neurons (Figure S5, Movie S6). We found ∼8% of β-actin mRNAs and RAVs co-localized/co-trafficked over time (Figure S5C). We next examined whether cell stimulation modifies RAV-mRNA interactions. cLTP did not significantly alter the frequency of these associations (cLTP: 13.9 ± 4.4%; pre-cLTP: 8.0 ± 3.2%; p = 0.5912) (Figure S5C). These data suggest that RAVs may similarly interact with β-actin mRNA under basal and stimulated conditions for local translation. Nevertheless, it remains to be seen whether this is specific to β-actin mRNA or is more broadly generalizable to other mRNAs.

Recent evidence suggests that the ER is involved in local translation within axons of primary neurons (*13, 27*). We therefore investigated whether local translation also occurs in dendrites using the puromycin-proximity ligation assay (Puro-PLA). Here, puromycin—acting as a tRNA analog—is incorporated into nascent polypeptides and subsequently detected using an anti-puromycin antibody. When combined with PLA, this method visualizes newly synthesized proteins at discrete sites of local translation (*43*). For the PLA component, we labeled endogenous Sec61β, a core element of the translocon complex in the ER. Puro-PLA showed puncta in MAP2+ dendrites, indicating putative local translation sites and suggesting that ER-associated translation occurs outside the cell body in dendrites (Figure S6). However, since Sec61β may be present in ER tubules and RAVs, it remained unclear if these puro-PLA puncta represented translation hotspots originating from one or both ER subcompartments. Therefore, to directly evaluate whether RAVs serve as platforms for local translation in distal neurites, we conducted single-molecule imaging of nascent peptides (SINAPS) via the SunTag reporter of translation alongside oScarlet-KDEL-labeled RAVs.

In response to cLTP stimulation, there was significantly increased association between SunTag signal representing ongoing translation and oScarlet-KDEL-labeled RAVs (Figure 4A, 4B, S7). We found a 2.3-fold increase in the percentage of SunTag puncta co-localized with RAVs 30 min after cLTP (cLTP: 38.2 ± 4.3%; basal: 16.6 ± 3.2%; p <0.0001) (Figure 4B). To confirm that SunTag signal reflects bona fide protein synthesis, we treated neurons with puromycin post-cLTP (100 µM, 30 min) to inhibit translation. Puromycin substantially attenuated SunTag puncta formation in both basal and cLTP-stimulated conditions along with its association with RAVs (cLTP + puromycin: 11.4 ± 2.3%; basal + puromycin: 15.8 ± 2.9%), validating the specificity and relevance of the SunTag assay to report local translational activity (Figure 4A, 4B, S7).

**Figure 4.**
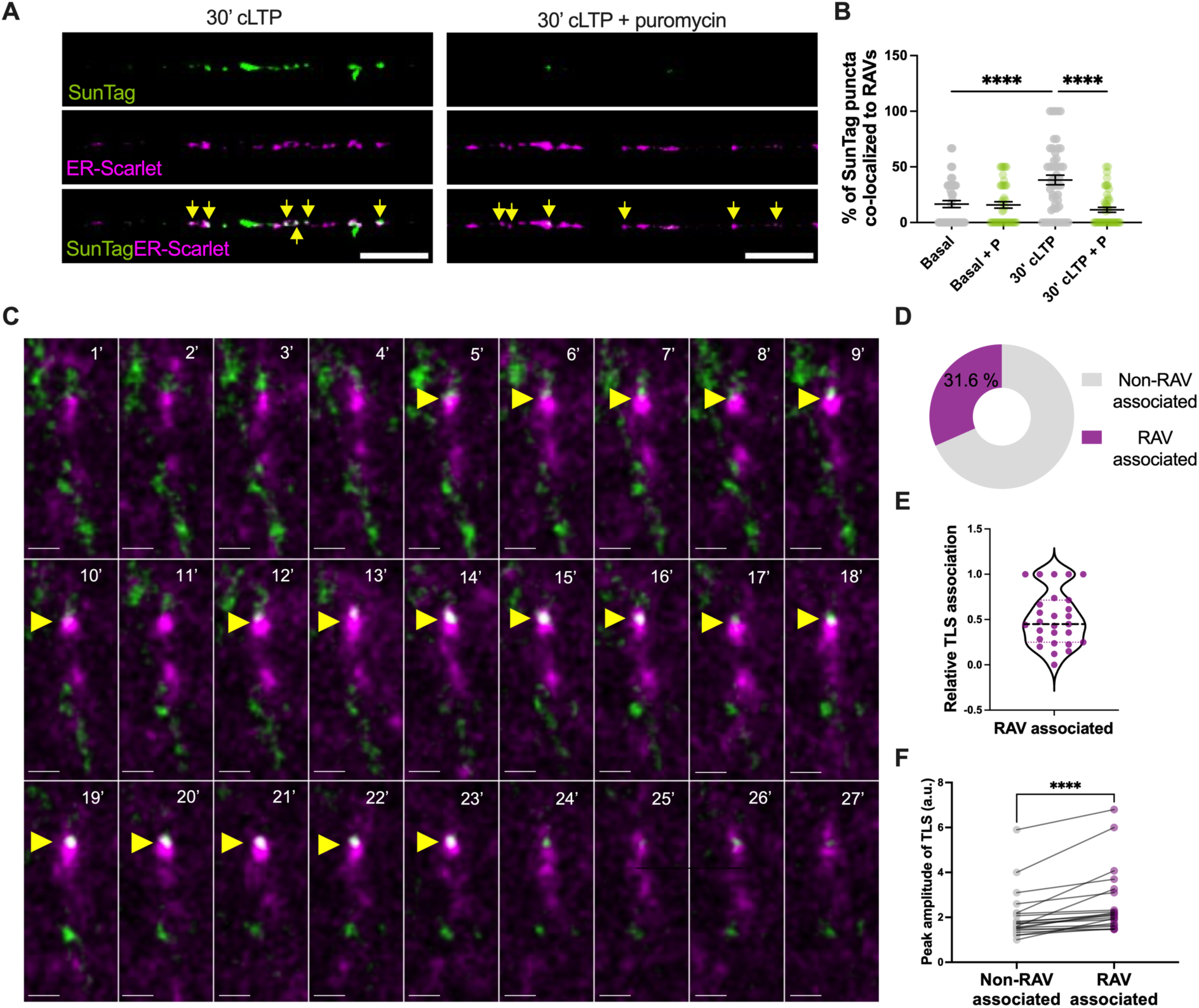
RAVs promote local activity-driven translation in dendrites. **(A)** Representative images of mouse primary cortical neurons co-expressing SunTag reporter (green) for labeling of translation sites (TLS) and oScarlet-KDEL (magenta) for RAV labeling in neurites. Immunolabelling of SunTag with anti-GCN4 alongside oScarlet-KDEL-labeled puncta (anti-KDEL) (yellow arrows) showed colocalization (in white) in response to cLTP stimulation (left panel). cLTP plus puromycin (P) treatment attenuated SunTag signal (right panel). Scale bar, 5µm. **(B)** Quantification of SunTag co-localization with oScarlet-KDEL labelled puncta. Two-way ANOVA with Tukey’s *post hoc* test. Mean ± SEM; ****p < 0.0001, n = 42, Basal; n = 44, Basal + P; n = 51, 30 min cLTP; n = 47, 30 min cLTP + P. **(C)** Time-lapse imaging of primary mouse hippocampal neurons expressing oScarlet-KDEL (magenta) and SunTag (green) reporter. In response to cLTP, there was increasing overlap (white) between oScarlet-KDEL-labeled RAVs and translation sites in dendritic segments. Scale bar, 1 µm. **(D)** Frequency of RAV-SunTag association across different dendrites, n = 19 dendrites. **(E**) RAV association with TLS relative to path trajectory. Each dot represents a translation site, n = 27. (**F)** Quantification of peak SunTag fluorescence intensity in the presence or absence of RAV association during the entire trajectory of translation. n = 21 trajectories. Wilcoxon test. ****p < 0.0001.

To determine how RAVs drive translation in response to cLTP, we performed live-cell imaging in hippocampal neurons expressing the SunTag reporter, scFv-sfGFP, and oScarlet-KDEL. Translation sites (TLS) were defined as discrete fluorescent foci brighter than diffusing proteins, resulting from the rapid binding of superfolder GFP-labeled single-chain antibody fragments (scFv) to the SunTag epitope as nascent peptides were synthesized. We found that stationary RAVs mostly colocalized with TLS in dendrites following cLTP (Figure 4C, Movie S7), indicating that RAVs actively translate mRNAs in response to neuronal stimulation. Quantitation showed that 31.6% of TLS were directly associated with RAVs (Figure 4D). Moreover, during the time course of translation, RAVs associated with TLS for 50.9 ± 0.1% of the sampling time (Figure 4E). Notably, RAVs often exhibited directed movement and accumulated near existing TLS, likely contributing to the ribosomal pool required for local translation (Figure S8, Movie S8). Indeed, tracking the integrated intensity of individual TLS associated with RAVs demonstrated greater peak SunTag amplitude (2.6 ± 0.3 a.u.) compared to non-RAV-associated TLS (2.0 ± 0.2 a.u.), suggesting that RAVs boost local translational output (Figure 4F). It is notable that translation also occurred independently of RAVs based on sites where SunTag signal did not directly overlap with RAVs. We posit that RAV-independent local protein synthesis may be carried out by other cellular components, including free ribosomes or other organelles such as endosomes that have been associated with translation (*11, 12, 15*). Nevertheless, the higher amplitude of RAV-associated TLS in response to neuronal activity suggests that RAVs boost translation in distal neuronal processes. Overall, our data point to RAVs as a key component of the machinery responsible for local translation in neurons.

## Discussion

Neurons rely on local protein synthesis (*1, 4, 8, 25, 26*), however, ribosomes are relatively sparse in the neurites, raising important questions about how ribosomes are trafficked to neurites to promote local protein synthesis. Classic models of ribosome trafficking relied primarily on diffusion and bulk flow which may be inadequate for the spatiotemporal precision required for activity-dependent plasticity (*1, 7*). Here, we suggest that RAVs offer a response to these long-standing questions by delivering ribosomes to specific sites in neurites in a directed, dynamic manner. Our findings establish RAVs as specialized platforms for the rapid, efficient delivery of ribosomes to sites requiring local protein synthesis. In neurites, RAVs are highly dynamic where changes in neuronal activity modify RAVs trafficking in a compartment-specific manner. Furthermore, RAVs associate with mRNAs to boost translation in spatially precise locations, thereby regulating activity-dependent protein synthesis in neurons. Together, these data provide a novel mechanism for local protein synthesis in neurons far from the cell body.

We extensively characterized the ultrastructural properties and dynamics of RAVs across different cellular systems utilizing a multi-scale imaging approach combining CLEM, cryo-ET, FIB-SEM VEM, and live-cell imaging. Cryo-CLEM, cryo-ET, and VEM data showed that RAVs are heterogenous throughout neurons. These structures differ in size, as well as in extent of ribosome association. While some RAVs are heavily decorated with ribosomes, others show sparser patterns of ribosome associations with RAV membranes. In some cases, cryo-CLEM data demonstrated vesicles that colocalize with an ER marker but lack significant ribosomal associations. These data therefore suggest that ribosomal associations with RAVs may be dynamic, offering neurons the flexibility required to adapt their translational machinery to changing demands for protein synthesis across different compartments. Just as importantly, these data add to the growing body of evidence showing that multiple membranous organelles (*e.g*., endosomes, lysosomes, mitochondria) can transiently associate with ribosomes (*11, 15*).

We showed that RAVs exhibit heterogeneous patterns of movement (*e.g*., stationary, processive, intermittent) in neurites. Moreover, microtubule depolymerization by nocodazole reduced the number of mobile RAVs, suggesting that RAVs rely on microtubule-based transport. This is consistent with previous findings demonstrating that depletion of the microtubule motor kinesin decreases both the number and speed of KDEL-positive vesicles (*34*). Studies examining mRNA trafficking have described similar dynamics, showing that specific mRNAs (*e.g*., *BDNF, Camk2a, Psd95, β-actin, Arc*) can be either mobile or stationary (*25, 40, 44-46*) and rely on microtubule motors for intracellular trafficking (*47, 48*). Moreover, these mRNAs become increasingly confined at local sites following neuronal stimulation where they may undergo local translation (*25, 40, 44*). The distinct patterns of RAV movement—stationary, processive, and intermittent—likely reflect differences in the mechanisms by which these organelles are trafficked and engage with mRNAs.

Strikingly, we identified compartment-specific changes in RAV dynamics in response to neuronal activity. The accumulation of stationary RAVs in dendrites following cLTP may provide a mechanism for rapidly increasing ribosomal capacity at specific sites on demand. This response matches the temporal demands for early-phase long-term potentiation (*21, 40, 49, 50*). In contrast, the acceleration of RAV transport in axons may ensure their efficient delivery to distant terminals without necessarily increasing local density. Overall, these data suggest that RAVs act differently in dendrites and axons, raising important questions about the molecular mechanisms driving compartment-specific differences.

Additional structures likely support RAVs in driving local translation across compartments and contexts. For example, axonal tubular ER can support local protein synthesis though the interaction of ribosomes and mRNAs with P180/RRBP1 (*13*). Endosomes similarly promote protein synthesis in axons (*11*). This raises many questions concerning the physiological relevance of ribosomes associated with different organelles versus those functioning freely. It is possible that free ribosomes handle constitutive translation of abundant proteins (*26*), while RAVs along with other ribosomes-associated organelles (*e.g*., endosomes, mitochondria) may drive translation in distinct functional niches of the neuronal arbor (*10, 51*). While ribosomes broadly drive protein synthesis, we posit that their localization and ability to function either in a free state or in association with different organelles may be critical for tight spatiotemporal regulation of protein synthesis.

A study limitation is the absence of a unique molecular marker for RAVs since RAVs represent a subcompartment of the ER. Therefore, it is difficult to distinguish RAVs from the larger ER network that can also participate in local translation. We relied on the ER retention signal, KDEL, to label RAVs, however, both the Golgi apparatus and ER-Golgi intermediate compartment (ERGIC) also contain KDEL receptors (*52, 53*). Thus, we cannot fully exclude the possibility that at least some of the observed KDEL-labeled puncta represent these organelles rather than pure RAVs. The resolution achieved with FIB-SEM VEM (∼5 nm) is one of the highest achievable with this technique (*54, 55*), but, remains insufficient to unambiguously resolve ribosomes or other fine structural details. Recent advances combining cryo-FIB milling in conjunction with cryo-ET in tissue provide the potential for high-resolution *in situ* visualization of ribosomes and macromolecular complexes within intact tissue environments under near-native conditions (*56-59*).

Overall, our study provides a mechanistic framework to better understand the regulation of local protein synthesis needed for synaptic plasticity. The rapid recruitment of RAVs to distal processes can deliver the ribosomal machinery necessary for local protein synthesis of plasticity-related proteins, including AMPA receptors, scaffolding proteins, and structural components (*7, 60, 61*). Several key questions emerge from our work that will guide future investigations: What are the molecular mechanisms controlling RAV biogenesis and whether RAVs exhibit functional specificity? Ultimately, this study sheds new light on how compartmentalized translation is orchestrated.

## ACKNOWLEDGEMENTS

We thank Ms. Mary Brady for expert assistance with movie generation. We are also grateful to Drs. James Olds, David Lewis, and Jamie Moy for technical assistance and helpful discussions.

## Funding

This work was supported by The Pittsburgh Foundation (ZF), the Commonwealth of Pennsylvania Formula Fund (ZF), the Baszucki Group (ZF), NIH R21MH122961 (SD), R01NS083085 (SD), R01NS122784 (MSG), and S10MH133575 (ZF, JRG). Cryo-ET studies were supported by NIH R35GM122588 (GJJ). This work was supported in part by the Koch Institute Support (core) Grant P30-CA14051 from the National Cancer Institute. We thank the Koch Institute’s Robert A. Swanson (1969) Biotechnology Center for technical support, specifically Peterson (1957) Nanotechnology Materials Core Facility (RRID:SCR_018674).

## Author contributions

Conceptualization: EMS, SDC, SD, ZF

Methodology: EMD, SDC, KK, GDN, DM, EKFD, JN, JRG, MAP, LE, CBM, KW, CMH

Investigation and data analysis: EMS, SDC, KK, GDN, JN, JRG, MAP, LH, LM, LE, PNJ, CBM, KW, CLM, ANF, SP, CMH, NK, DM, EKFD, JLO, KB, GJJ, SCW, KD, LEF, MSG, ZPW, AKRLJ, SD, ZF

Visualization: EMS, SDC, KK, GDN, JN, JRG, MAS, PNJ, CBM, KW, SD

Writing – original draft: EMS, SDC, SD, ZF

Writing – review & editing: EMS, SDC, KK, SD, ZF

## Competing interests

ZF is the recipient of an investigator-initiated grant from UPMC Enterprises. The remaining authors declare that they have no competing interests.

## Data and materials availability

All data are available in the main text or the supplementary materials. Correspondence and requests for materials should be addressed to ZF and SD.

## Supplementary Materials

Materials and Methods

Supplementary References

Figures S1 to S8

Tables S1 to S3 Movies S1 to S8

## Notes

### Summary of Updates

We include a new section on room temperature correlative light and electron microscopy data, revised volume electron microscopy data, updated author names and affiliations, and updated supplemental materials..

