## Supplementary Materials for "Ribosome-Associated Vesicles promote activity-dependent local translation in neurons"

**This PDF includes:**

Materials and Methods

Supplementary References

Figures S1 to S8

Tables S1 to S3

**Other Supplementary Material for this manuscript includes the following:**

Movies S1 to S8

### Materials and Methods

**Chemicals.** Chemicals used in the present study were purchased from Sigma-Aldrich (St. Louis, MO) and Tocris Bioscience (Bristol, UK), unless otherwise specified.

**DNA constructs.** mNeon-KDEL was constructed via addition of a leader sequence from mouse immunoglobulin κ light chain 5′ to mNeonGreen followed by a KDEL ER retrieval sequence 3′ to the fluorophore as described previously (*1*). oScarlet-KDEL was generated from the mNeon-KDEL construct by substituting mNeonGreen with codon-optimized mScarlet-I (oScarlet). ER-mScarletI (Addgene #137805) consists of mScarlet-I fused to the KDEL sequence 3′ to the fluorophore (*2*). NLS-stdMCP-stdGFP consists of a fusion of synonymous versions of MCP (stdMCP) and GFP (stdGFP) with a nuclear localization sequence (NLS) (Addgene #98916). A fluorescent SunTag reporter of mRNA translation was modified from the original SINAPS construct (pUbC-FLAG-24xSuntagV4-oxEBFP-AID-baUTR1-24xMS2V5-Wpre; Addgene #84561) (*3*). The reporter consists of the mouse *β-actin* 3´ UTR sequence based on a design from the laboratory of Dr. Robert Singer (Albert Einstein College of Medicine, Bronx, NY). pHR-scFv-GCN4-sfGFP-GB1-NLS-dWPRE encodes an antibody that binds to the SunTag GCN4 peptide which is also fused to sfGFP and contains an NLS sequence (Addgene #60906).

#### Viral packaging

**AAV molecular cloning and production.** The mNeon-KDEL coding sequence was subcloned into an AAV vector backbone consisting of AAV2 ITR sequences, a human elongation factor 1α (EF1α) promoter, a woodchuck hepatitis virus posttranscriptional regulatory element (WPRE), and a human growth hormone polyadenylation sequence (hGH polyA) to generate AAV-EF1α-mNeon-KDEL-WPRE-polyA. Chimeric AAV was produced as previously described (*4*). Briefly, HEK293FT cells were quadruple transfected with AAV-EF1α-mNeon-KDEL-WPRE-polyA,

AAV2/1 rep/cap, AAV 2/2 rep/cap, and pDF6. After 48 hours, cells were harvested, lysed, and the supernatant was applied to a heparinized column. The column was washed with NaCl solution, followed by viral elution and concentration. Purity was determined by denaturing SDS-PAGE gel electrophoresis and function was validated in primary rat hippocampal neurons.

**Lentiviral molecular cloning and production.** Constructs were cloned into the phage-UbC-RIG lentiviral vector backbone, enabling expression of the respective inserts under the control of the UbC promoter. Packaging of the constructs into lentiviral particles was conducted as previously reported (*5, 6*). Briefly, lentivirus was generated by mixing plasmids for ENV (pMD2.VSVG), packaging (pMDLg/pRRE), and REV (pRSV-Rev). Human embryonic kidney (HEK) 293T cells were subsequently transfected with expression vectors for oScarlet-KDEL, NLS-stdMCP-stdGFP, scFv-sfGFP, or SunTag construct along with the lentiviral packaging plasmids using calcium phosphate. Virus-containing supernatant was concentrated using the Lenti-X concentrator (Clontech Laboratories, Mountain View, CA), resuspended in Neurobasal A medium, and stored at -80°C until transduction. A subset of the scFv-sfGFP and SunTag constructs were also commercially packaged into lentivirus by PackGene Biotech (Houston, TX).

**Animal husbandry**

Animals were housed and handled in accordance with NIH and institutional guidelines. We abided by ARRIVE guidelines for reporting animal research (*7, 8*). Mice and rats were housed in cages with a 12:12 light:dark cycle and always had access to food and water *ad lib*. All studies were approved by the Institutional Animal Care and Use Committees at the University of Pittsburgh, Albert Einstein College of Medicine, and Emory University. Every effort was made to ameliorate animal suffering.

**Tissue culture**

**Primary rat neuron culture and sample preparation.** Dissociated rat cortical and hippocampal neurons were isolated from Long–Evans rats (Charles River Laboratories, Wilmington, MA) of both sexes at embryonic day 18 (E18) as described earlier (*1, 9*). Neurons were maintained in neurobasal medium supplemented with B27 (100 units/mL), penicillin/streptomycin (100 mg/mL), and 2 mM glutamine with two-fifths of the media replaced every 4 days (37°C, 5% $CO_2$).

For live-cell imaging studies, neurons were plated onto 35 mm-diameter circular glass-bottom MatTek dishes (MatTek, Ashland, MA) coated with poly-D-lysine (20 µg/mL) and laminin (3.4 µg/mL) at a density of 70,000-80,000 cells per MatTek dish. Neurons were transfected on days 8-10 *in vitro* (DIV) using Lipofectamine 2000 (Invitrogen, Waltham, MA) with a total of 1-2 µg/well of DNA. The transfected neurons were imaged at DIV10-13.

For cryo-correlative light and electron microscopy (cryo-CLEM) and cryo-electron tomography (cryo-ET) studies, freshly isolated neurons were plated onto sterile, poly-D-lysine/laminin-coated 200 mesh Quantifoil R2/2 gold NH2 London finder grids (200 mesh) (Quantifoil/SPT Labtech, Covina, CA) at a density of $4\text{-}6\times10^4$ cells/mL as previously described (*10, 11*). Neuron-coated grids were maintained in 35 mm glass bottom MatTeK dishes (37°C, 5% $CO_2$). Neurons were transfected on DIV10-14 with mNeon-KDEL using Lipofectamine 2000. 24 hours after transfection, neurons were plunge-frozen in liquid ethane/propane using a Vitrobot Mark IV (Thermo Fisher Scientific-FEI, Hillsboro, OR) and stored in liquid nitrogen until imaging. Just prior to plunge-freezing, a fluorescent microsphere/gold solution was applied to grids as reported earlier (*1*). 500 nm blue polystyrene fluoropheres (Phosphorex Inc., Hopkinton, MA) were diluted in phosphate-buffered saline (PBS) and mixed with 20 nm colloidal gold (Sigma-Aldrich) pretreated with bovine serum albumin. The fluorospheres and 20 nm colloidal gold both served as fiducial markers to facilitate the registration of cryogenic fluorescence microscopy images collected from different channels as well as with cryo-EM projection images for cryo-CLEM (*1*).

**Primary mouse hippocampal neuron culture and sample preparation.** Mouse hippocampal neurons were isolated as described previously (*12*). Hippocampi were isolated from homozygous Actb-MBS knock-in (Actin$^{M/M}$) mice (*13*); neurons from both sexes were combined in cultures. Hippocampal tissue was digested in 0.25% trypsin (37°C, 15 min), triturated, and plated onto poly-D-lysine-coated MatTek dishes and 12 mm glass coverslips at a density of 75,000 neurons/glass coverslip for subsequent live-cell imaging and immunocytochemistry studies. Neurons were cultured in Neurobasal A medium supplemented with B-27, GlutaMAX, and primocin (InvivoGen, San Diego, CA).

For mRNA trafficking studies, neurons were transduced with NLS-stdMCP-stdGFP lentivirus at DIV8-10, followed by imaging at DIV14-15; a subset of experiments employed neurons transduced at DIV4-5 and imaged at DIV10.

For SunTag experiments, neurons were co-transduced at DIV5-6 with oScarlet-KDEL, SunTag reporter, and ScFv-sfGFP to concurrently monitor RAV dynamics alongside mRNA translation in peripheral neurites, with imaging conducted at DIV15-16.

**Primary mouse cortical neuron culture and sample preparation.** Cortical neuron were cultured from mouse E17 embryos (WT C57BL/6J strain #027, Charles River lab as previously described (*14*). Briefly, mouse cortices were digested in 0.25% trypsin without EDTA (Thermo Fisher Scientific) for 10 min and then plated in 10% FBS containing neurobasal (Gibco) for 2 hours. Neurons were then switched to neurobasal media with B27 supplement (Gibco) and fed twice weekly.

**Primary mouse dorsal root ganglion culture and sample preparation.** Dorsal root ganglion (DRG) neurons were isolated from 18-week-old male CGRP$^{CreER/+}$ mice (*15*) (kindly provided by Dr. Jami Saloman, University of Pittsburgh, Pittsburgh, PA). Cre recombinase was induced three weeks prior by four consecutive daily intraperitoneal injections of 75mg/kg tamoxifen (dissolved in corn oil). Mice were deeply anesthetized with isoflurane (0.3 ml/L, 60-90 sec) and perfused with ice-cold PBS before L3-L5 DRGs were rapidly dissected, enzymatically treated, and mechanically dissociated as previously described (*16*). Dissociated neurons were transduced with AAV-EF1α-mNeon-KDEL-WPRE-polyA and placed directly into compartmentalized microfluidic devices (DOC540, Xona Microfluidics, Research Triangle Park, NC) in Neurobasal medium supplemented with 0.1% penicillin/streptomycin (Gibco), B27 (Gibco), 10 ng/mL NGF (Sigma-Millipore), and 10 ng/mL GDNF (R&D Systems, Minneapolis, MN) at 37°C in 5% $CO_2$. Media was changed every other day, and cells were analyzed on DIV6–10.

Glass and PDMS devices were cleaned with 70% ethanol, air-dried for 1 hour, and bonded using oxygen plasma activation (Harrick Plasma, Inc., Ithaca, NY). Devices were then coated within 10 minutes with 5 µg/mL mouse laminin (Gibco BRL, Gaithersburg, MD, USA) and 0.1 mg/mL poly-L-ornithine. To guide the terminals through the microchannels, the terminal chambers were flooded with 50 ng/mL NGF and 50 ng/mL GDNF to generate a growth factor gradient.

**Live-cell imaging**

**Imaging cultured rat primary neurons and DRG neurons.** Time-lapse wide-field imaging of cultured cortical and DRG neurons was performed on a Nikon TiE microscope (Nikon Corp., Tokyo, Japan) equipped with a Plan Apo VC 60× Oil DIC N2/1.4 NA oil immersion objective (Olympus, Tokyo, Japan) and a Prime 95B Scientific CMOS (sCMOS) camera (Teledyne Photometrics, Tucson, AZ). 488-nm, 561-nm, and 640-nm lasers were used for illumination along with corresponding dichroic filters. Samples were imaged in a temperature-controlled Tokai Hit microscope stage top incubator (Amuza Inc., San Diego, CA) set to 37°C with 5% $CO_2$. Images were collected as single planes across time at 100 msec exposure times over 1-5 min using Nikon NIS-Elements software. The field-of-view was 1200 pixels × 1200 pixels, with a pixel size of 0.183 μm. The camera was set to 1×1 binning with 16-bit digitalization.

**Imaging cultured mouse primary hippocampal and cortical neurons.** Time-lapse imaging of cultured mouse hippocampal neurons was conducted on a wide-field fluorescence microscope consisting of an Olympus IX-81 base equipped with an iXon Ultra DU-897U EMCCD camera (Andor Technology Ltd, Belfast, UK), and UPlanApo TIRF 150x 1.45 NA oil immersion objective (Olympus). 491-nm (Calypso-25, Cobolt, San Jose, CA), 561-nm (LASOS-561-50, Lasertechnik GmbH, Germany), and 640-nm (CUBE 640-40C, Coherent Inc, Santa Clara, CA) lasers were delivered through the back port with laser power controlled via an acousto-optic tunable filter (AOTFnC-400.650-TN, AA Opto-Electronic, Orsay, France). Lasers were reflected by a 4-band excitation dichroic filter (Di01-R405/488/561/635; Semrock, Rochester, NY). Emission filters (FF01-525/50 for green, FF01-605/64 for red; Semrock) were mounted on a motorized filter wheel (FW-1000, Applied Scientific Instrumentation) for rapid switching between wavelengths. Samples were imaged in a temperature-controlled Tokai Hit microscope stage top incubator (Amuza) set to 37°C with 5% $CO_2$. Axons and dendrites were distinguished based on their morphological characteristics as well as via immunolabeling for axonal versus dendritic markers. Time-lapse single plane images of the targeted sites were acquired at 800 msec time intervals (1.2 Hz) under basal unstimulated and stimulated conditions.

Immunofluorescence images of fixed neurons were acquired on an inverted Nikon Ti2 epifluorescence microscope using a 60× oil immersion objective (Plan Apo, NA 1.42). Samples were illuminated with a SOLA light engine (Lumencor), and fluorescence emission was detected using an ultra-low readout noise, back-thinned sCMOS camera (ORCA-Fusion BT, Hamamatsu Photonics). Z-stacks spanning 4 µm total depth were acquired with 0.2 µm optical sections using NIS-Elements Advanced Research software (version 6.20.01, Nikon).

**Drug treatments**

**Neuronal stimulation.** Primary cultured neurons were stimulated via chemical long-term potentiation (cLTP) as described previously (*5, 17*). Briefly, cultured neurons were pre-incubated with 50 µM APV (2-amino-5-phosphonovaleric acid) overnight (12-14 hours), followed by cLTP induction via treatment with 100 µM picrotoxin and 200 µM glycine in $Mg^{2+}$-free Hibernate A medium (BrainBits, Springfield, IL) for 5 min. Neurons were subsequently washed twice and returned to Hibernate A medium containing $Mg^{2+}$ and $Ca^{2+}$ for live-cell hippocampal neuron imaging studies; cortical neurons were imaged in complete Neurobasal A medium. For experiments testing protein synthesis-dependent association of RAVs with the SunTag reporter, neurons were incubated immediately after cLTP stimulation with 100 µM puromycin hydrochloride (Millipore Sigma).

**Microtubule depolymerization.** Primary hippocampal neuron microtubules were depolymerized by exposure to 1 µM nocodazole for 2 hours. Time-lapse images were acquired as described above, the resulting images analyzed using the KymoTool plugin in the ImageJ software package (National Institutes of Health, Bethesda, MD) (*18*).

**Puromycin proximity ligation assay (Puro-PLA).** Puro-PLA was performed as reported earlier (*19*) with additional modifications. Briefly, rat hippocampal neurons (DIV 12-15) were incubated with puromycin (3 µM) for 10 min prior to fixation. As controls, neurons were either incubated in the absence of puromycin or pretreated with anisomycin (40 µM) for 30 min before the addition of puromycin. Neurons were subsequently fixed with 4% paraformaldehyde (PFA)/4% sucrose for

10 min at room temperature, washed 3x with PBS, and permeabilized with 0.1% Triton X-100 in PBS 1x for 15 min. The cells were next incubated with blocking solution provided in the NaveniFlex Cell Red *in situ* proximity ligation assay (PLA) kit (Navinci Diagnostics, Uppsala, Sweden). Prior to commencing PLA, the fixed neurons were immunostained with polyclonal guinea pig anti-MAP2 antibody (Synaptic Systems GmbH, Göttingen, Germany; #188004, RRID:AB_2138181). Following overnight incubation with primary antibody (1:1000 dilution in NaveniFlex antibody diluent) at 4°C, cells were washed 3x for 10 min with PBS and then incubated with Alexa Fluor 647 goat anti-guinea pig secondary antibody (Thermo Fisher Scientific; #A21450, RRID:AB_141882) for 1 hour at room temperature. Neurons were subsequently immunostained overnight with the following antibodies: monoclonal mouse anti-puromycin (clone 12D10, Sigma-Aldrich; #MABE343, RRID:AB_2566826) and rabbit polyclonal anti-Sec61B (Proteintech, Rosemont, IL; #51020-2-AP, RRID:AB_2186413); antibodies were diluted in NaveniFlex antibody diluent (anti-puromycin: 1:500; anti-Sec61B: 1:100). The PLA assay was performed according to manufacturer's recommendations. Neurons were mounted in ProLong$^{TM}$ Gold antifade reagent (Invitrogen/Thermo Fisher Scientific, Carlsbad, CA). Samples were imaged on a Nikon Crest spinning disk confocal microscope equipped with a 40× oil objective (Nikon Plan Fluor 40x oil DIC H/N2, NA 1.3). Images were processed with ImageJ software.

**Immunocytochemistry**

Primary mouse cortical neurons were plated on 12 mm glass coverslips at a density of 75,000 cells/coverslip. Neurons were transduced with oScarlet-KDEL and/or SINAPs (SunTag reporter) lentivirus at DIV7. Following 5 days of expression, cells were preincubated overnight with 50 µM APV and cLTP-stimulated the next day. Cells were then fixed with 4% PFA/4% sucrose for 10 min. After washing 3× with PBS, neurons were permeabilized with 0.1% Triton X-100 (in PBS) for 15 min at room temperature, followed by blocking with 10% BSA (1 hour, room temperature).

For experiments to assess differences in RAV localization in axons and dendrites, neurons were immunostained with the following primary antibodies at room temperature for 90 min: rabbit anti-mCherry to label mScarlet-ER (Abcam, Waltham, MA; #ab167453, RRID:AB_2571870, 1:500), mouse anti-NEFL-M for axon labeling (Developmental Studies Hybridoma Bank, Iowa City, IA; #2H3, RRID:AB_531793, 1:50), and guinea pig anti-MAP2 for dendrite labeling (Synaptic Systems; #188004, RRID:AB_2138181, 1:2500). Primary antibodies were diluted in 2% BSA (in PBS). Samples were washed 3× for 5 min with PBS, followed by incubation with Alexa Fluor-conjugated secondary antibodies for 30 min at room temperature: donkey anti-rabbit IgG Alexa Fluor 568 (Thermo Fisher Scientific; #A10042, RRID:AB_2534017, 1:1000), goat anti-mouse IgG Alexa Fluor 647 (Thermo Fisher Scientific; #A-21235, RRID:AB_2535804, 1:1000), and donkey AffiniPure anti-guinea pig IgG Alexa Fluor 488 (Jackson ImmunoResearch, West Grove, PA; #706-545-148, RRID:AB_2340472, 1:1000).

For experiments to evaluate RAV colocalization with translation sites, neurons were immunostained with rabbit anti-mCherry antibody to label mScarlet-ER (Abcam; #ab167453, 1:500) and mouse anti-GCN4 antibody to label SunTag (Absolute Antibody, Newark, CA; clone C11L34, #Ab00436-1.4, 1:250). Samples were then incubated with donkey anti-rabbit IgG Alexa Fluor 568 (Thermo Fisher Scientific; #A10042, RRID:AB_2534017, 1:1000) and goat anti-mouse IgG Alexa Fluor 647 (Thermo Fisher Scientific; #A-21235, 1:1000).

Cells were washed 3× for 5 min each with PBS and mounted in ProLong Diamond Antifade Mountant (Thermo Fisher Scientific, P36930). Cells were imaged on an inverted Nikon Ti2 epifluorescence microscope using a 60× oil immersion objective (Plan Apo, NA 1.42). Samples were illuminated with a SOLA light engine (Lumencor), and fluorescence emission was detected using an ORCA-Fusion BT sCMOS camera (Hamamatsu Photonics). Z-stacks spanning 4 μm total depth were acquired with 0.2 μm optical sections using NIS-Elements Advanced Research software (version 6.20.01, Nikon). Images were maximum intensity projected for analysis.

**Live-cell imaging data analysis**

**RAV trafficking*.*** Raw images were deconvolved via the Richardson-Lucy algorithm in NIS-Elements software with a limit of 10 iterations. No other processing was performed on the images. Subsequently, RAV dynamics were examined in distal neurites via semi-automated spot detection algorithms employed by the Imaris software package (Oxford Instruments, Zurich, Switzerland). Results were independently validated in ImageJ software using the Kymolyzer plugin (*20*) which quantified: the percentage of RAVs in motion, percentage of stationary RAVs, RAV speed and displacement. Stationary RAVs were defined as immobile if they remained 100% still throughout imaging. Mobile RAVs were defined according to periods of movement during imaging including processive, continuous movement (total movement = 100% of imaging time) or intermittent movement (total movement <100% of imaging time).

**RAVs and mRNA interactions.** To establish interactions between β-actin mRNA with RAVs, we used the KymoTool plugin in ImageJ. Initially, kymographs representing both β-actin mRNA and RAVs were merged and visually inspected to identify sites of overlap. Upon identification of potential colocalization events, RAV-mRNA distance was calculated using the formula $d=\sqrt{((x_b - x_a)^2 + (y_b - y_a)^2)}$, based on the provided point coordinates. Colocalization was established when the calculated distance was < 0.5 μm for at least four consecutive frames.

**RAV co-localization at translation sites.** Co-imaging of SunTag signal and RAVs on the same dendrite was performed to identify RAV co-localization at translation sites (TLS). To reduce potential background signal from the SunTag reporter, a time-averaged (3 frame) time series was used for the analysis. To be considered a TLS, the intensity needed to be 20% higher than background scFv-sfGFP signal. For a TLS to be RAV-associated, an overlap for at least 3 frames within a 500 nm distance was used as an inclusion criterion. Semi-automated tracking of TLS was performed in ImageJ software using the Trackmate plugin (*21*) and a previously described custom script in MATLAB software (MathWorks, Natick, MA)(*3, 5*); a gap of 3 frames was treated as the same site. Since RAVs often associated with TLS for a specific duration, the fraction of total TLS trajectory length that showed RAV-association was quantified (relative TLS association). The intensity of an individual TLS, which is a measure of the amount of nascent peptide synthesis, was recorded for the entire trajectory in the presence or absence of RAV-association (peak amplitude).

**Immunocytochemistry data analysis.** For quantification of RAV localization to dendrites and axons, immunofluorescence images were analyzed using the Analyze Particles macro in the Fiji image analysis software package (ImageJ, NIH). Images were projected according to maximum intensity, and dendritic or axonal segments (labeled with MAP2 or NEFL-M, respectively) were straightened using Fiji's segmented line tool. Fluorescence intensity thresholds were maintained constant across experimental groups. RAVs were identified by size thresholding (50–800 nm diameter). For analysis of RAV colocalization with SunTag-labeled sites, dendrites were straightened in Fiji, and independent thresholds were applied to SunTag and RAV channels. Each SunTag-labeled punctum was evaluated for colocalization with RAV puncta or clusters within the same dendritic segment. Following established protocols (*5*), SunTag signals that could not be resolved as discrete puncta were excluded. Distal dendritic segments were preferentially analyzed due to sparser RAV and SunTag labeling to facilitate unambiguous determination of colocalization events.

**Cryo-fluorescence microscopy and cryo-electron tomography.** Plunge-frozen EM grids were clipped in auto-grids (Thermo Scientific) and neurons were imaged using a Nikon CFI S Plan

Fluor 60× NA 0.7 extra-long-working-distance air-objective (WD 2.62–1.8 mm) as previously described (*22*). mNeon-KDEL was visualized with a GFPHQ filter set [Semrock; Excitation (Ex) 455-485 nm, DM 495 nm, Emission (Em) 500-545 nm]. Blue fluorospheres were visualized with a DAPI filter set (Semrock; Ex 340-380 nm, DM 400 nm, Em 435-48 5nm). Red autofluorescence was imaged using a TxRed filter set (Semrock; Ex 540-580 nm, DM 595 nm, Em 600-660 nm). Regions of interest containing mNeon-KDEL-positive puncta were identified using cryo-LM microscopy. These regions were correlated and targeted for tilt-series collection with a low-magnification EM image using previously described cryo-CLEM methods (*1, 23*). Cryo-EM grids previously imaged by cryo-light microscopy (cryo-LM) were subsequently imaged by cryo-ET using Titan Krios transmission electron microscopes (Thermo Fisher) equipped with K2 or K3 cameras (Gatan, Inc., Pleasanton, CA) at 300 keV.

**Volume reconstruction, subtomogram extraction, alignment, and averaging.** Each tilt series was collected with the Volta Phase Plate bi-directionally from -60° to +60° in 2° increments using SerialEM software (*24*) (University of Colorado Boulder, Boulder, CO) at 3-6 μm underfocus. A total of 100-140 electrons/$Å^2$ was distributed evenly across the tilt-series. The image pixel size used for subtomogram averaging was 2.650 Å (unbinned). Subsequent subtomogram averaging was performed via the EMAN2 software tomography pipeline (*25*). Initially, unbinned tilt-series were automatically aligned and reconstructed using EMAN2. In total, 5 cryo-tomograms were generated to provide sufficient particles for further processing. 164 particles were picked using EMAN2 particle picking software with a box size of 36 × 36 × 36 pixels (unbinned = 144 pixels). An initial model was then generated in C1 with all 164 particles in 3 iterations. This model was used for subtomogram averaging refinement. The fourth iteration map was employed as a model in a second round of subtomogram averaging refinement using a keep value of 0.8. The particles were mapped back into the cryo-tomogram using EMAN2 software.

### Room temperature correlative light and electron microscopy

**Fixation and sample preparation.** All reagents were freshly prepared on the experimental day. Primary rat cortical neurons were initially fixed in 2% PFA (Electron Microscopy Sciences, Hatfield, PA; #EMS 15700) (in culture media) for 30 min at room temperature followed by subsequent fixation in 4% PFA (in 0.1M PIPES) for 24 hours at 4°C. After fixation, cells were washed 3× in 0.1 M PIPES and incubated for 10 minutes in 0.15% glycine. Cells were infiltrated with a thin layer of 12% pig skin gelatin for 30 min at 37°C and solidified on ice for at least 30 min before being infiltrated with 2.3 M sucrose for 48 hours at 4°C under gentle shaking. Small blocks (~0.5 $mm^3$) were cut from the infiltrated, solidified gelatin, mounted with the cells facing up on sample pins for cryo-ultramicrotomy (Electron Microscopy Sciences, #EMS 75959-06), and frozen in liquid nitrogen where they were stored until sectioning.

**Cryo-ultramicrotomy.** Sectioning was performed using a Leica UC7 cryo-ultramicrotome (Leica, Wetzlar, Germany) equipped with a cryo-diamond knife (DiATOME, Fort Washington, PA; cryo-immuno 35° diamond knife). Semithin sections (500 nm) were cut at -80°C and collected onto 12 mm glass coverslips using a mixture of equal parts of 2.3 M sucrose and 2% methylcellulose (25 centipoises). Sections were stored at room temperature until further processing.

**Immunostaining and fluorescence microscopy.** For immunofluorescence, coverslips with cryosections were incubated in 0.1 M PIPES at 37°C for 30 min to remove the sucrose-methylcellulose layer. Samples were then blocked in 1% BSA-c (Aurion Biotech, Seattle, WA; #900.099) in 0.1M PIPES for 30 min at room temperature before being incubated with Alexa Fluor 488-conjugated anti-KDEL antibody (Abcam; #EPR12668, RRID:AB_2819147); the antibody was diluted 1:100 in blocking buffer. Finally, coverslips were washed 3× in 0.1 M PIPES, followed by a final wash in $ddH_2O$, and mounted onto glass slides using DAPI-containing mounting medium

(Invitrogen, #D3571) for nuclear staining. The samples were imaged on a Zeiss LSM 880 confocal microscope equipped with a 63× oil objective with 1.4 NA (Zeiss, Oberkochen, Germany).

**Electron microscopy and image correlation.** After fluorescence imaging, slides were unmounted and stained with heavy metal for electron microscopy. Briefly, the samples were exposed to reduced osmium (2% osmium tetroxide, 2.5% potassium ferrocyanide) for 15 min, washed 3× in 0.1M PIPES buffer, and then stained again with an aqueous solution consisting of 1% tannic and reduced osmium in a ratio of 60:1, respectively. Samples were then washed 3× in water, stained for 15 min in 1% uranyl acetate (in water), washed again 3× in water before undergoing serial dehydration in increasing ethanol concentrations (50%, 70%, 90%, 100%), followed by resin infiltration. All steps were performed on ice, with ice-cold media. Infiltrated samples were baked for at least 48 hours at 60°C before being trimmed and sectioned with a Leica UC7 ultramicrotome. A diamond knife (DiATOME) cut ultrathin sections (60 nm), which were then collected in carbon-coated formvar slot grids (Electron Microscopy Services, #FCF2010-Cu-EA) and imaged on a Tecnai T12 transmission electron microscope (FEI). Image registration was performed using BigWarp, a Fiji plugin for landmark-based registration of large image datasets (*26*). To ensure unbiased registration, landmarks were selected based on prominent cellular features visible (*e.g.*, nucleus, axons, vacuoles) in both brightfield and electron microscopy images.

**Focused ion beam-scanning electron microscope volume electron microscopy (FIB-SEM VEM)**

**Human subject.** The brain specimen was obtained during a routine autopsy conducted at the Allegheny County Office of the Medical Examiner (Pittsburgh, PA) after consent for donation was obtained from next-of-kin. An independent committee of experienced research clinicians confirmed the absence of any lifetime psychiatric or neurologic diagnoses for the decedent based on medical records, neuropathology examinations, toxicology reports, as well as structured diagnostic interviews conducted with family members of the decedent (*27*). A subject with demographic and tissue features within the range of our previously published light and electron microscopic studies of postmortem human brain tissue (*28-31*) was selected. This subject was a 62-year-old male who died suddenly and out-of-hospital with an accidental manner of death. The postmortem interval (defined as the time elapsed between death and brain tissue preservation) was 6.0 hours. Brain tissue pH was measured as 7.0. All procedures were approved by the University of Pittsburgh's Committee for the Oversight of Research and Clinical Training Involving Decedents and the Institutional Review Board for Biomedical Research.

**Human brain tissue preparation and imaging.** All procedures have been previously described in detail (*32*). Briefly, a fresh tissue block dissected from the middle frontal gyrus Brodmann Area 46 was fixed in 4% paraformaldehyde and 0.2% glutaraldehyde for 24 hours at room temperature, followed by 24 hours at 4°C in fresh fixative. Vibratome sections (50 µm) were stained and resin-embedded using a previously reported approach (*33*) with minor modifications (*32*), including dehydration steps completed at room temperature, the use of propylene oxide during the dehydration steps and a different resin mixture containing EMbed 812, Araldite GY 502, DDSA (Dodecenyl Succinic Anhydride) and BMDA (N=Benzyl-N, N-Dimethylamine)(Electron Microscopy Sciences). After infiltration with 100% resin, tissue sections were set at 60°C for 48 hours to allow for resin polymerization. After polymerization was complete, a subsample of cortical layer 3 was dissected and adhered to a resin capsule. Excess resin was trimmed via a Leica Ultracut UCT ultramicrotome equipped with a diamond knife (DiATOME) until the face of the tissue was visible. The tissue sample was then imaged via FIB-SEM VEM using a Helios 5 CX DualBeam FIB-SEM (Thermo Fisher Scientific, Hillsboro NanoPort), as described in detail earlier (*32*). Specifically, the sample was milled using a gallium ion beam with a 5 nm slice thickness and SEM-imaged using 2 keV landing energy, 400 pA beam current, and 4 µs dwell time. Individual digital images were aligned and registered via Amira software (Thermo Fisher Scientific) to generate the complete 3D volume. This volume was denoised, and structures of interest were segmented with Amira software.

**Statistical analyses**

Statistical significance was determined using GraphPad Prism (version 10, GraphPad Software, Inc., La Jolla, CA). Statistical analyses used in this study include Two-tailed t-test, Mann-Whitney test or non-parametric one-way ANOVA, (*i.e.*, Kruskal-Wallis test and Friedman test) and two-way ANOVA. Statistical significance is defined as $*p < 0.05$, $**p < 0.01$, $***p < 0.001$, $****p < 0.0001$.

| | Pre-cLTP | cLTP (15-25 min) | cLTP (60-70 min) |
|---|---|---|---|
| **Neurites analyzed** | 37 | 37 | 37 |
| **# Total RAVs** | 185 | 253 | 220 |
| **RAVs/μm** | 0.022 ± 0.0025 | 0.032 ± 0.0032 | 0.031 ± 0.0043 |
| **Average Speed (μm/s)** | 0.76 ± 0.062 | 0.72 ± 0.052 | 0.65 ± 0.048 |

**Table S1. RAV analyses in rat cortical neurons in response to cLTP.** All values represent mean ± SEM, n = 3 independent experiments.

| | Pre-cLTP | cLTP (15-25 min) |
|---|---|---|
| **Neurites analyzed** | 95 | 118 |
| **# Total RAVs** | 222 | 340 |
| **RAVs/µm** | 0.071 ± 0.005 | 0.085 ± 0.005 |
| **Average Speed (µm/s)** | 0.69 ± 0.04 | 0.86 ± 0.05 |

**Table S2. RAV analyses in mouse hippocampal neurons in response to cLTP.** All values represent mean ± SEM, n = 3 independent experiments.

| | Axons | | Dendrites | |
|---|---|---|---|---|
| | **Pre-cLTP** | **cLTP:15'** | **Pre-cLTP** | **cLTP:15'** |
| **Neurites analyzed** | 66 | 87 | 29 | 31 |
| **# Total RAVs** | 155 | 221 | 67 | 119 |
| **# Stationary RAVs** | 37 | 70 | 44 | 78 |
| **# Moving RAVs** | 118 | 151 | 23 | 41 |
| **RAVs/µm** | 0.074 ± 0.061 | 0.080 ± 0.006 | 0.062 ± 0.005 | 0.099 ± 0.011 |
| **Stationary RAVs/µm** | 0.017 ± 0.004 | 0.023± 0.004 | 0.040 ± 0.005 | 0.067± 0.012 |
| **Moving RAVs/µm** | 0.056 ± 0.050 | 0.057 ± 0.006 | 0.022 ± 0.006 | 0.032 ± 0.084 |
| **Average Speed (µm/s)** | 0.70 ± 0.04 | 0.86 ± 0.05 | 0.65 ± 0.10 | 0.84 ± 0.09 |

**Table S3. RAV analyses in mouse hippocampal axons and dendrites in response to cLTP.** All values represent mean ± SEM, n = 3 independent experiments.

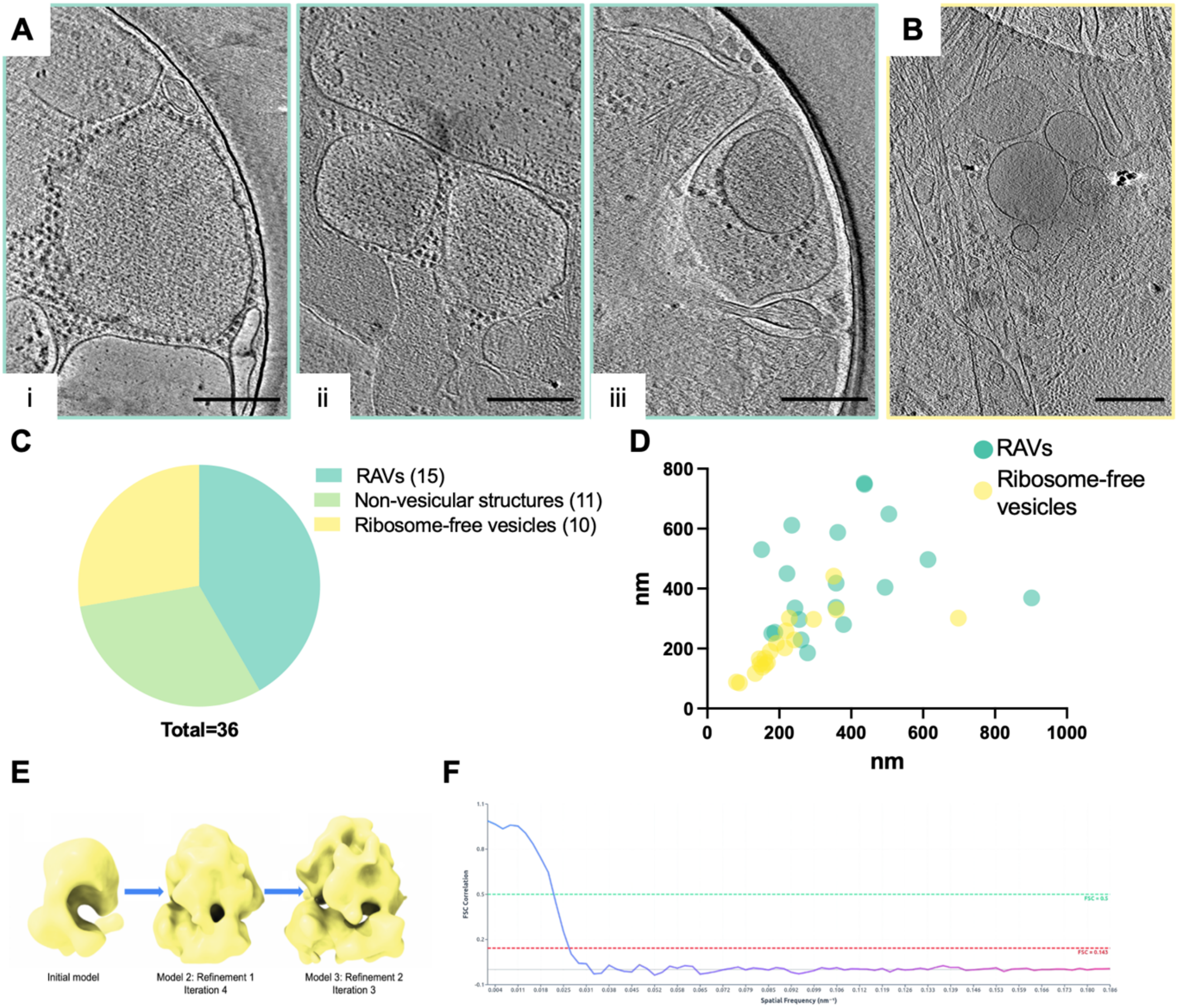


**Figure S1. Examples of KDEL-colocalizing vesicles in primary neurons via cryo-CLEM. (A)** Representative cryo-tomographic slices showing multiple RAVs with membrane-associated ribosomes in hippocampal primary neurons; these samples are from a cryo-CLEM dataset and showed co-localization with mNeon-KDEL fluorescence. **(i)** Well-decorated RAV showing numerous membrane-associated ribosomes. **(ii, iii)** RAVs partially decorated with ribosomes. Scale bars, 250 nm. **(B)** Cryo–tomographic slice featuring ribosome-free vesicles that also showed co-localization with mNeon-KDEL fluorescence. Scale bars, 250 nm. **(C)** Quantification of cryo-tomograms featuring mNeon-KDEL-positive structures (n=36 tomograms). **(D)** Size distribution of RAVs versus ribosome-free vesicles (RAVs, n=19, ribosome-free vesicles, n=18). **(E)** Model generation for alignment and subtomogram averaging of *in situ* RAV-bound ribosomes in neurons. We first generated a model from a total of 164 particles in 3 iterations. Then, 4 iterations of subtomogram averaging generated a second model for further subtomogram averaging and refinement. A third model was subsequently generated during the second round of refinement (iteration 3), demonstrating clear features of an 80S ribosome. **(F)** Fourier Shell Correlation (FSC) curve to estimate the final resolution of the ribosome subtomogram average. The FSC=0.143 and 0.5 criterion (red dashed line and green dashed line respectively) were used. The final map resolution is estimated at 37.9 Å (FSC=0.143).

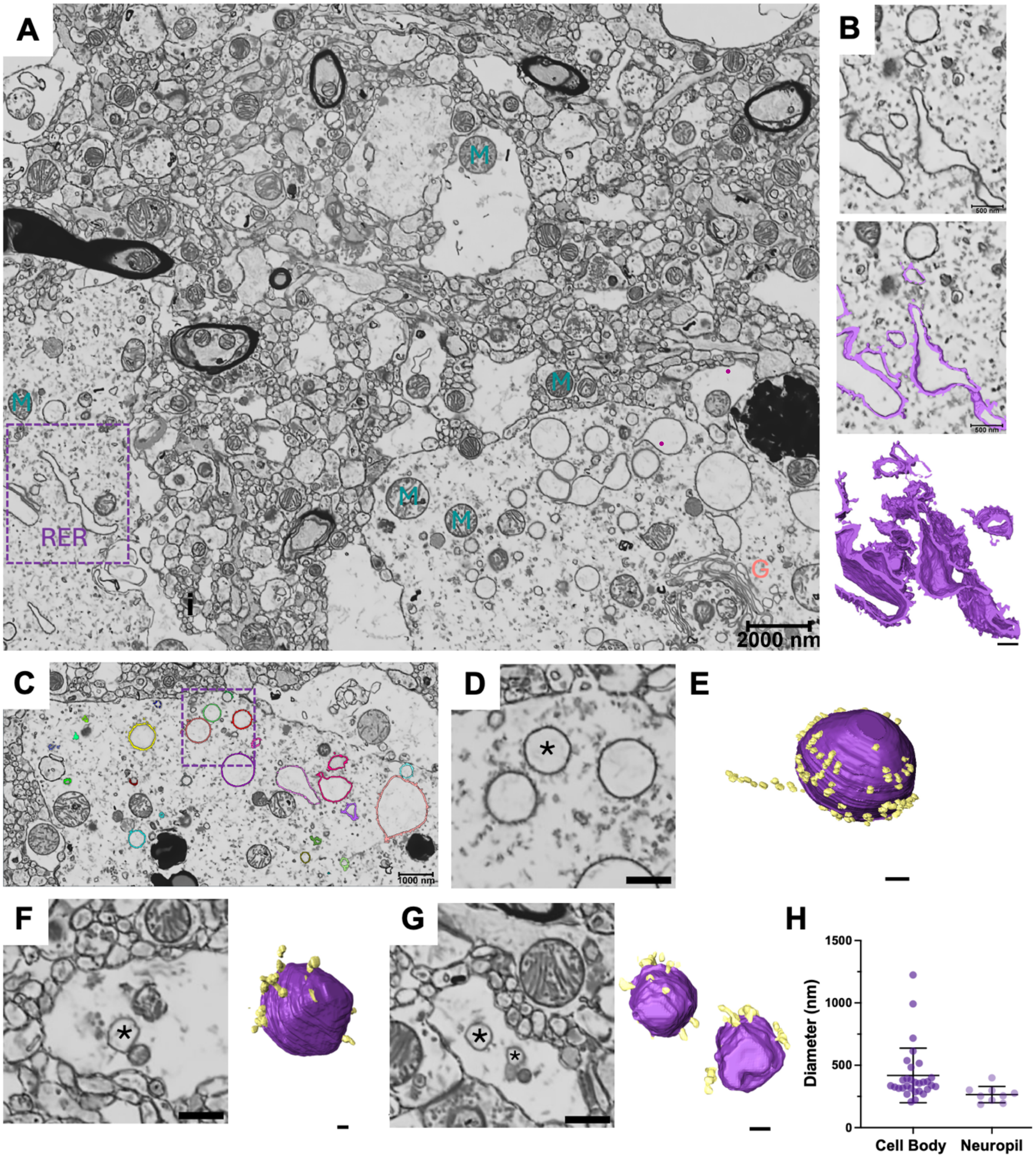


**Figure S2. Focused ion beam-scanning electron microscopy (FIB-SEM) reveals RAVs in human cortical brain tissue. (A)** A representative FIB-SEM image illustrating excellent preservation of brain tissue subcellular architecture. This includes numerous organelles distributed throughout the imaged area including the Golgi apparatus (G), mitochondria (M), and rough endoplasmic reticulum (RER) network. Scale bar, 2 μm. **(B)** An enlarged view of the boxed area in Panel A shows intact RER network (i), its segmentation (ii), and 3D reconstruction (iii). Scale bar, 500 nm **(C)** SEM image highlighting RAVs within the cell body. Scale bar, 1000 nm **(D)** An enlarged view of the boxed region in Panel C of a representative RAV. Scale bar, 500 nm. **(E)** 3D reconstruction of the RAV in Panel D (*) shows ribosome-like particles (in yellow) associated with the RAV membrane (in purple). Scale bar, 200 nm. **(F-G)** FIB-SEM images of the neuropil featuring RAVs alongside their respective 3D reconstructions. Scale bar, 500 nm SEM image, 100 nm 3D reconstruction. Asterisks denote RAVs. **(H)** Quantification of the RAV vesicle diameter in the cell body and neuropil. Mean ± SEM (n = 30, cell body; n = 9, neuropil).

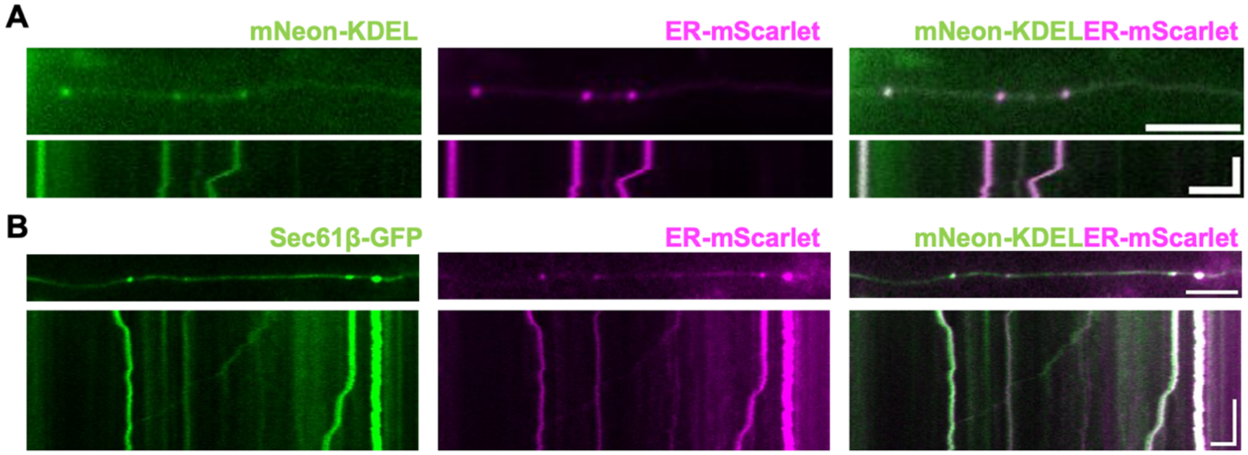


**Figure S3. Different ER markers label dynamic puncta in primary neurons. (A)** Colocalization of mNeon-KDEL and ER-Scarlet-KDEL in rat hippocampal neurons. **(B)** Colocalization of Sec61β and ER-Scarlet-KDEL in rat hippocampal neurons. For each panel: the top row shows a snapshot from a time-lapse sequence in a neurite. Scale bar, 10 μm. Bottom row shows the corresponding kymograph, illustrating the dynamic behavior of ER-labeled puncta.  Scale bar, 5 μm, 20 sec.

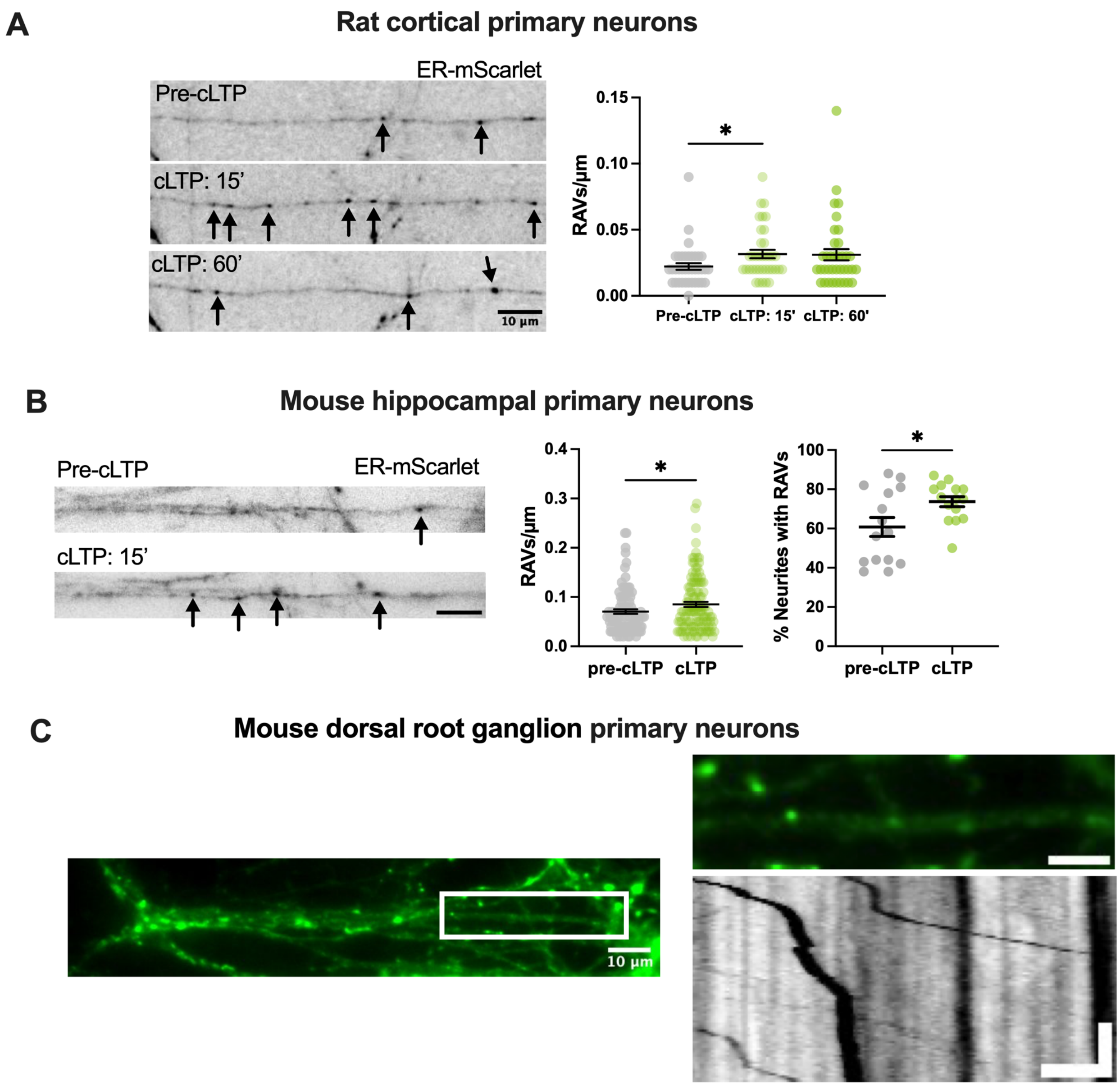


**Figure S4. RAV trafficking in different neuron types. (A)** Representative images showing increased local recruitment and accumulation of RAVs (indicated by arrows) within cortical neurites in response to chemically induced long-term potentiation (cLTP) (left panel). Quantification of RAV density (right panel). Scale bars, 10 µm. Mean ± SEM; *$p<0.05$, Friedman test. **(B)** Representative images showing increased local recruitment and accumulation of RAVs (indicated by arrows) within hippocampal neurites in response to cLTP (left panel). Quantification of RAV density (middle panel), and percentage of RAV+ neurites (right panel). Scale bars, 5 µm. Mean ± SEM; *$p<0.05$, unpaired t-test. **(C)** Representative image of mNeon-KDEL-labeled RAVs in axons of primary mouse dorsal root ganglion neurons (left panel; scale bar, 10 µm). Magnified region highlighting dynamic RAVs (see white box, left panel) alongside a representative kymograph of a moving RAV (bottom panel; scale bar, 5 µm, 20 sec).

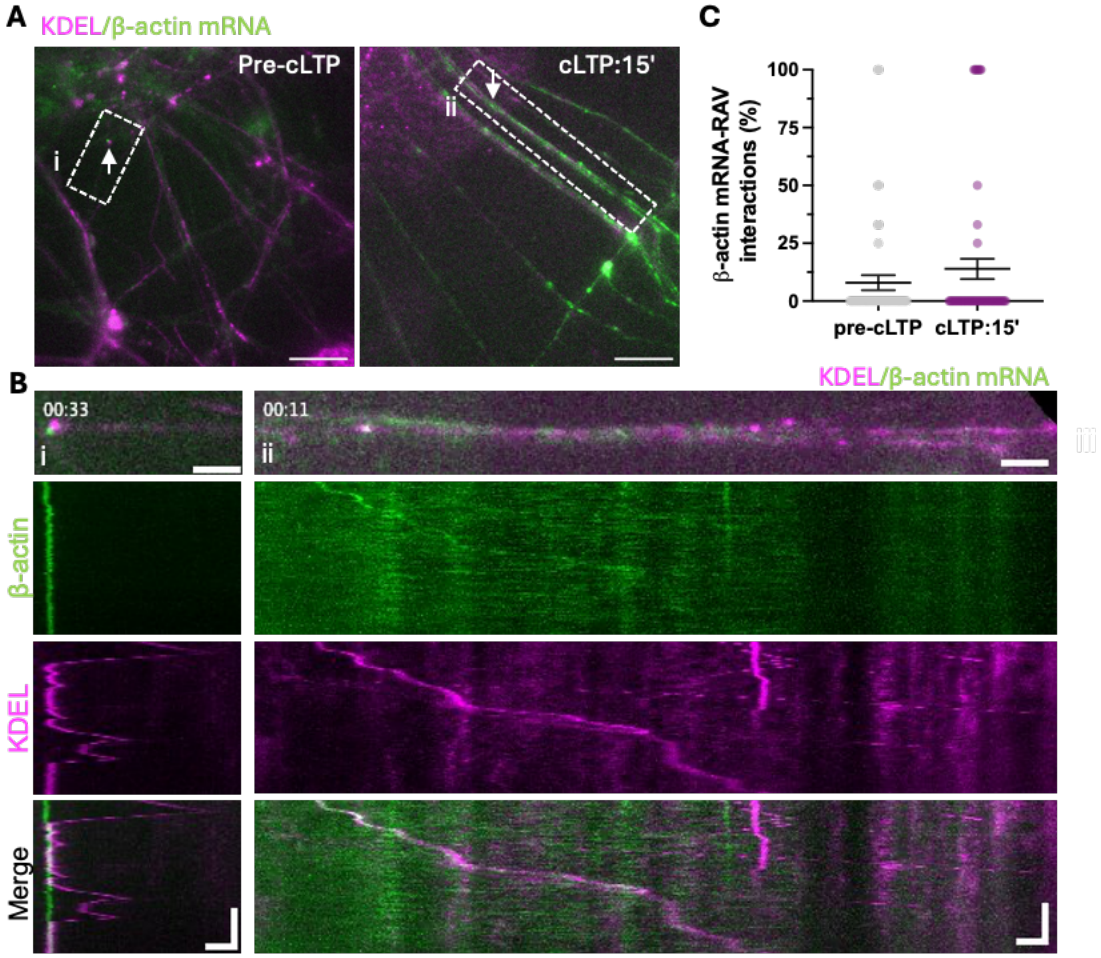


**Figure S5. RAVs associate with β-actin mRNA. (A)** Representative images of primary mouse hippocampal neurons expressing oScarlet-KDEL (magenta) and β-actin mRNA (green) under pre-cLTP and cLTP-stimulated (15 min) states. RAV-mRNA associations in neurites are highlighted with arrows. Scale bar, 20 µm **(B)** Enlarged view of neurites featured in boxes i and ii from Panel A with accompanying kymographs showing RAV-mRNA associations. Scale bars, 2 µm, 20 sec. **(C)** Quantification of β-actin mRNA-RAV interactions. pre-cLTP: n = 49; cTLP: n = 58. Mean ± SEM, Mann-Whitney test.

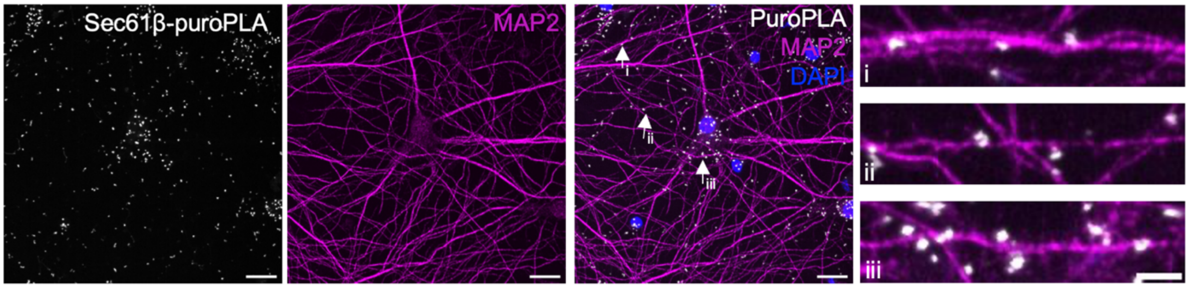


**Figure S6. Puromycin-proximity ligation assay (Puro-PLA) assay labels local ER translation sites in dendrites**. Imaging of primary rat hippocampal neurons previously treated with puromycin (3 μM, 10 min) prior to fixation. Puro-PLA was performed using anti-puromycin and anti-Sec61β antibodies to detect nascent polypeptides in close proximity to the ER translocon. Dendrites were labeled by MAP2 (magenta) and nuclei via DAPI stain (blue). Scale bars, 20 μm. Arrows indicate discrete local translation sites in dendrites which are shown at higher magnification in the insets (i, ii, iii). Scale bar, 5 μm.

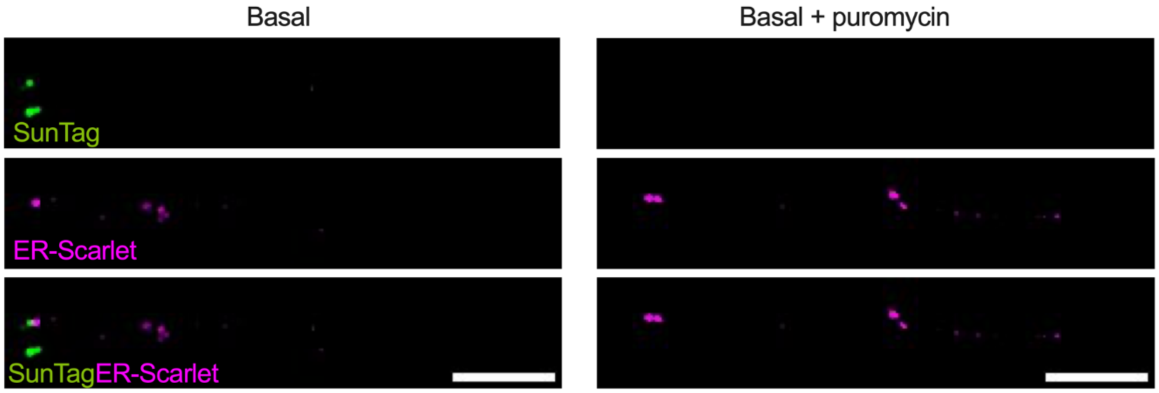


**Figure S7. SunTag reporter and oScarlet-KDEL co-expression in cortical neurons under basal conditions.** Representative immunofluorescence images of mouse primary cortical neurons co-expressing SunTag reporter (green) and oScarlet-KDEL (magenta). Immunolabelling of SunTag with anti-GCN4 alongside oScarlet-KDEL-labeled puncta (anti-KDEL) under basal (left panels) and basal + puromycin (right panels) conditions. Scale bars, 5 μm.

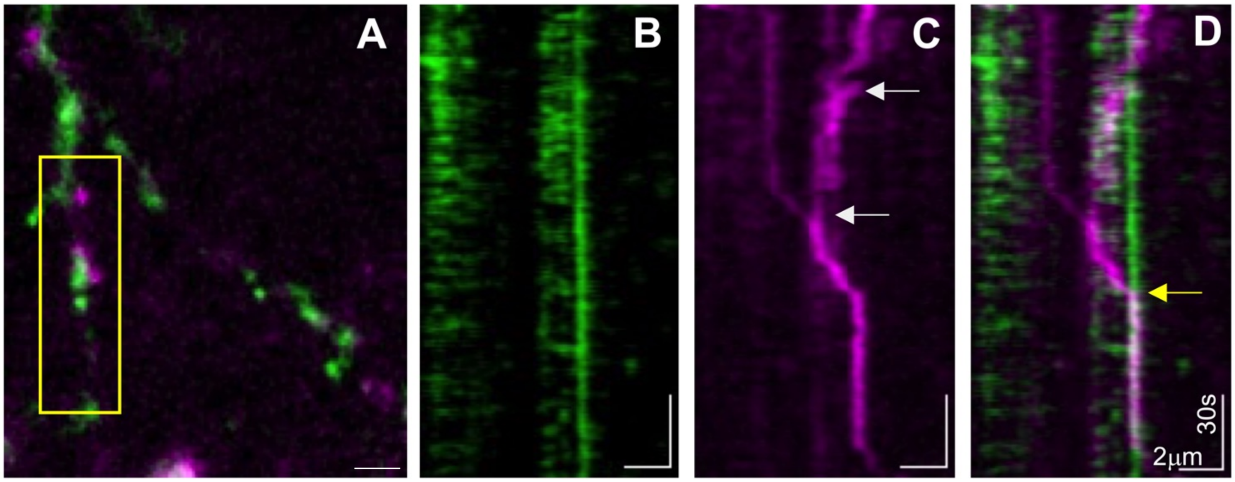


**Figure S8. RAVs cluster and accumulate at local translation sites. (A)** Dendritic region of a mouse primary hippocampal neuron co-expressing oScarlet-KDEL (magenta) and the SunTag translation reporter (green). Scale bar, 2 μm. **(B-D)** Kymographs generated from the selected dendritic region in Panel A (yellow box). Individual kymographs of SunTag **(B**, green**)** and oScarlet-KDEL-labeled RAVs **(C**, magenta**)** are shown. White arrows indicate points of RAV clustering during the imaging time course. **(D)** Merged kymograph indicates accumulation of RAVs at translation sites (yellow arrow). Scale bars: 2 μm, 30s.

**Supplementary Movies**

**Movie S1.** RAV in primary neurons visualized via cryo-ET. Scale bar, 200 nm.

**Movie S2.** RAV in human cortical brain tissue visualized via FIB-SEM VEM.

**Movie S3.** Mitochondria and endoplasmic reticulum are preserved in human cortical brain tissue imaged via FIB-SEM VEM.

**Movie S4.** RAV trafficking dynamics in primary cortical neurons via live-cell imaging.

**Movie S5.** RAVs traffic in axons of DRG neurons. White arrows indicate moving RAVs.

**Movie S6.** RAVs (magenta) transiently associate with β-actin mRNA (green).

**Movie S7.** RAV-driven local translation in response to neuronal stimulation. Yellow arrowhead indicates RAV association with SunTag reporter. Scale bar, 1 μm.

**Movie S8.** RAVs cluster at translation sites. Yellow arrow indicates points of RAV accumulation. Scale bar, 2 μm
